# Fate Before Function: Specification of the Hair Follicle Niche Occurs Prior to its Formation and Is Progenitor Dependent

**DOI:** 10.1101/414839

**Authors:** Ka-Wai Mok, Nivedita Saxena, Nicholas Heitman, Laura Grisanti, Devika Srivastava, Mauro Muraro, Tina Jacob, Rachel Sennett, Zichen Wang, Yutao Su, Lu M. Yang, Avi Ma’ayan, David M. Ornitz, Maria Kasper, Michael Rendl

**Affiliations:** Black Family Stem Cell Institute, Icahn School of Medicine at Mount Sinai, Atran Building AB7-10C, Box 1020; 1428 Madison Ave, New York, NY 10029, USA; Department of Cell, Developmental and Regenerative Biology, Icahn School of Medicine at Mount Sinai, Atran Building AB7-10C, Box 1020; 1428 Madison Ave, New York, NY 10029, USA; Graduate School of Biomedical Sciences; Icahn School of Medicine at Mount Sinai, Atran Building AB7-10C, Box 1020; 1428 Madison Ave, New York, NY 10029, USA; Oncode Institute, Hubrecht Institute–KNAW (Royal Netherlands Academy of Arts and Sciences) and University Medical Center Utrecht, 3584 CT Utrecht, the Netherlands; Department of Biosciences and Nutrition and Center for Innovative Medicine, Karolinska Institutet. 141 83 Huddinge, Sweden; Department of Pharmacological Sciences, Mount Sinai Center for Bioinformatics, BD2K-LINCS Data Coordination and Integration Center, Knowledge Management Center for Illuminating the Druggable Genome (KMC-IDG), Icahn School of Medicine at Mount Sinai, One Gustave L. Levy Place, New York, NY 10029, USA; Department of Developmental Biology, Washington University School of Medicine, St. Louis, Missouri, 63110, USA; Department of Dermatology, Icahn School of Medicine at Mount Sinai, Atran Building AB7-10C, Box 1020; 1428 Madison Ave, New York, NY 10029, USA

**Keywords:** stem cell niche, dermal condensate precursors, dermal papilla, hair follicle morphogenesis, hair follicle formation, cell fate specification, placode progenitors, single cell

## Abstract

Cell fate transitions are essential for specialization of stem cells and their niches, but the precise timing and sequence of molecular events during embryonic development are largely unknown. Here, we show that dermal condensates (DC), signaling niches for epithelial progenitors in hair placodes, are specified before niche formation and function. With 3D/4D microscopy we identify unclustered DC precursors. With population-based and single-cell transcriptomics we define a molecular time-lapse of dynamic niche signatures and the developmental trajectory as the DC lineage emerges from fibroblasts. Co-expression of downregulated fibroblast and upregulated DC genes in niche precursors reveals a transitory molecular state following a proliferation shutdown. Waves of transcription factor and signaling molecule expression then consolidate DC niche formation. Finally, ablation of epidermal Wnt signaling and placode-derived FGF20 demonstrates their requirement for DC-precursor specification. These findings uncover a progenitor-dependent niche precursor fate and the transitory molecular events controlling niche formation and function.

**Graphical Abstract:** 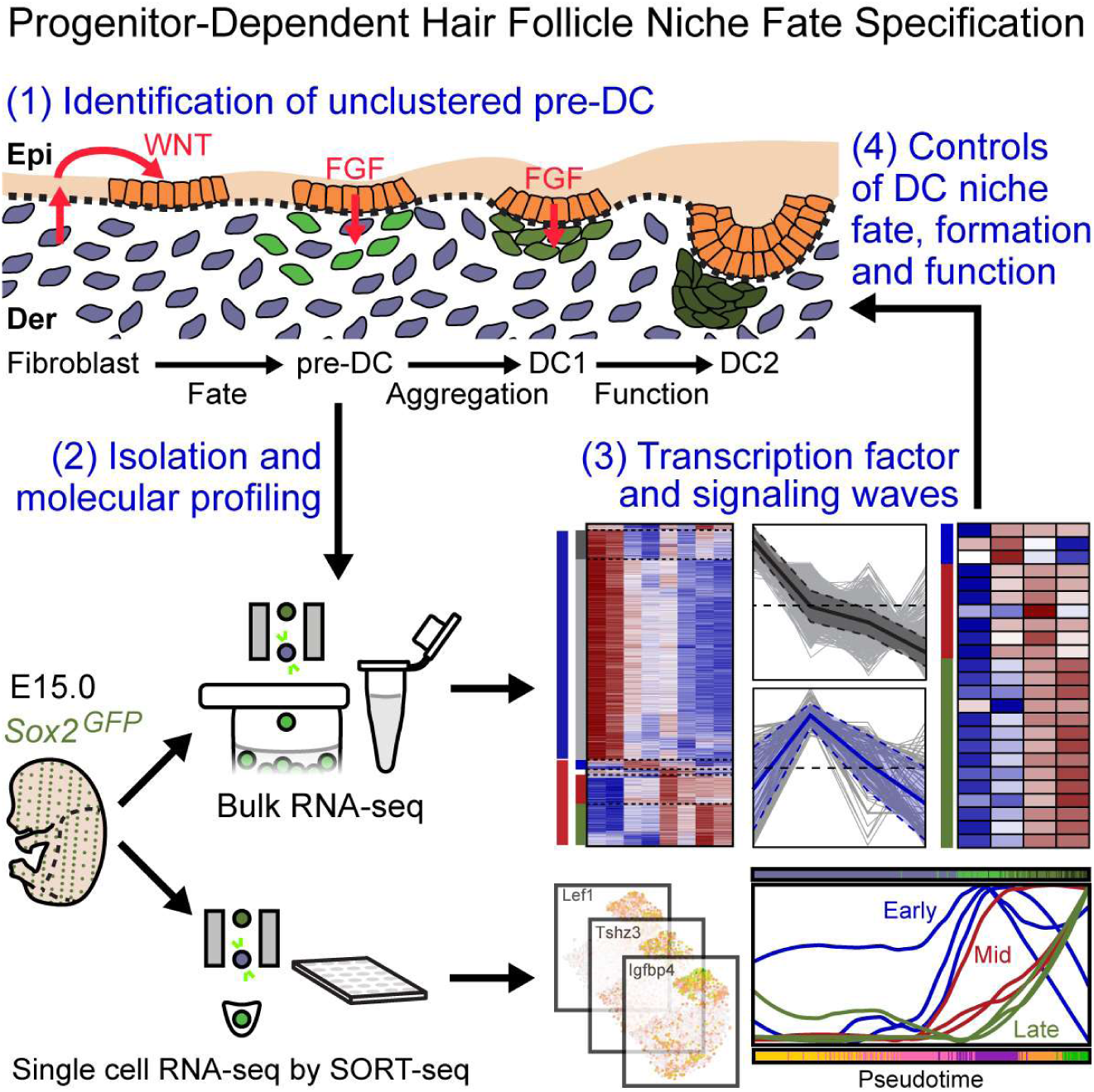

**HIGHLIGHTS:**

- Precursors of the hair follicle niche are specified before niche cluster formation
- Bulk/single cell RNA-seq defines early niche fate at molecular transitional state
- Successive waves of transcription factor/signaling genes mark niche fate acquisition
- Niche fate acquisition is not “pre-programmed” and requires FGF20 from progenitors

## INTRODUCTION

Many adult tissues are maintained during homeostasis or repaired after injury by resident stem cells as a resource of reserve cells with multilineage potential. Stem cell functions are either intrinsically controlled or are typically regulated by a large variety of neuronal, hormonal and other inputs from the microenvironment or niche (Chacón-Martínez et al., 2018; Ge and Fuchs, 2018; Heitman et al., 2018; Morrison and Spradling, 2008; Rezza et al., 2014) In many cases, signals are derived from specialized niche cells that control stem cell quiescence for long periods of time, regulate stem cell self-renewal, or guide their differentiation into one or more lineages of the respective tissue. Although neighboring niche cell types and their signals have been identified in several tissues such as bone marrow (Morrison et al., 2014; Tamma and Ribatti, 2017; Wei and Frenette, 2018), intestine (Shoshkes-Carmel et al., 2018), brain (Katsimpardi and Lledo, 2018), lung (Lee et al., 2017) and skin (Gonzales and Fuchs, 2017; Hsu et al., 2014b; Rompolas and Greco, 2014; Sennett and Rendl, 2012), our understanding of the molecular controls of niche fate specification remains limited. Equally unclear is the precise timing with which niche cells emerge during embryonic development relative to stem/progenitor cell fates and their mutual impact on each other.

In mice, hematopoietic stem cells first emerge from the dorsal aorta, then home to the placenta, spleen, and liver during successive developmental stages before finally relocating to the bone marrow (Dzierzak and Bigas, 2018). There periarteriolar stromal cells (Kunisaki et al., 2013) and osteoblasts (Calvi et al., 2003; Zhang et al., 2003) provide niche signals for quiescent hematopoietic stem cells and multipotent progenitors (Morrison et al., 2014; Wei and Frenette, 2018) for continued blood production, but when and how the niche is specified remains unknown. Similarly, specialized glial cells in the neurogenic niche (Falk and Götz, 2017), perialveolar fibroblasts close to lung progenitors (Nabhan et al., 2018), and interstitial fibroblasts surrounding the crypt stem cells in the intestine (Stzepourginski et al., 2017) have been identified as rich sources of niche signals to regulate nearby stem and progenitor cell functions in their respective systems. How these specialized niche cells acquire their stem cell supporting fate and how this relates to stem cell specification during development is largely unexplored.

In hair follicles, rapidly dividing progenitors in the proximal bulb region receive signals from the centrally-located dermal papilla (DP) niche, which controls hair follicle progenitor proliferation, migration and differentiation into several lineages of the hair shaft and its channel during continuous hair growth (Avigad Laron et al., 2018; Clavel et al., 2012; Morgan, 2015; Sennett and Rendl, 2012; Yang et al., 2017). During the hair cycle after bulb degeneration and rest, the DP niche sends activating signals to the stem cells in the hair germ to regenerate a new hair bulb (Hsu et al., 2014a). Many studies have investigated the developmental processes and morphogenetic events during embryonic hair follicle formation, including the emergence of stem cell and niche compartments (Chen et al., 2012; Millar, 2002; Ouspenskaia et al., 2016; Sennett and Rendl, 2012; Zhang et al., 2008, 2009). Around embryonic day (E) 12.5, secreted epidermal Wnts activate broad dermal Wnt/β-catenin signaling (Chen et al., 2012; Zhang et al., 2009). This is upstream of still unidentified dermal signals that at ∼E13.5 induce the fate specification of hair follicle progenitors in patterned pre-placodes (**Figure 1A,** hair follicle stage 0) (Paus et al., 1999) that can be molecularly identified by *Dkk4* and *Edar* expression (Sick et al., 2006; Zhang et al., 2009). Progenitor cell rearrangement, compaction, and migration then form the physically identifiable placode (Pc), likely due to continued dermal signals (**Figure 1A,** stage 1) (Ahtiainen et al., 2014). During this process, Pc progenitors signal back to the dermis for formation of dermal condensates (DC) (**Figure 1A,** stage 1, DC1), fibroblast-derived specialized cell clusters that act as signaling niches for placode progenitors to regulate continued hair follicle development (Millar, 2002; Sennett and Rendl, 2012; Sennett et al., 2015). Pc progenitors also give rise to suprabasal Sox9^+^ precursors of the future adult hair follicle stem cells in the bulge after completion of hair follicle morphogenesis (Ouspenskaia et al., 2016; Xu et al., 2015). Continued signal exchange between the DC and Pc leads to progenitor proliferation and downgrowth of the polarized hair germ (**Figure 1A,** stage 2, DC2), with the DC at the leading edge that, after an elongated hair peg stage, becomes engulfed by the progenitors to form the DP niche of the hair-shaft producing bulb (Grisanti et al., 2013; Sennett and Rendl, 2012). Several key signaling pathways have been identified for progenitor and niche formation (Biggs and Mikkola, 2014; Millar, 2002; Sennett and Rendl, 2012). Wnt/β-catenin signaling is essential and the most upstream for placode (Huelsken et al., 2001; Zhang et al., 2009) and condensate (Tsai et al., 2014) formation. Downstream Eda/Edar signaling is required for maintaining Wnt signaling and placode stabilization (Schmidt-Ullrich, 2006; Zhang et al., 2009). FGF20, another Wnt target, is a key placode-derived signal required for DC niche formation (Huh et al., 2013) in a process involving cell aggregation (Biggs et al., 2018).

**Figure 1.**
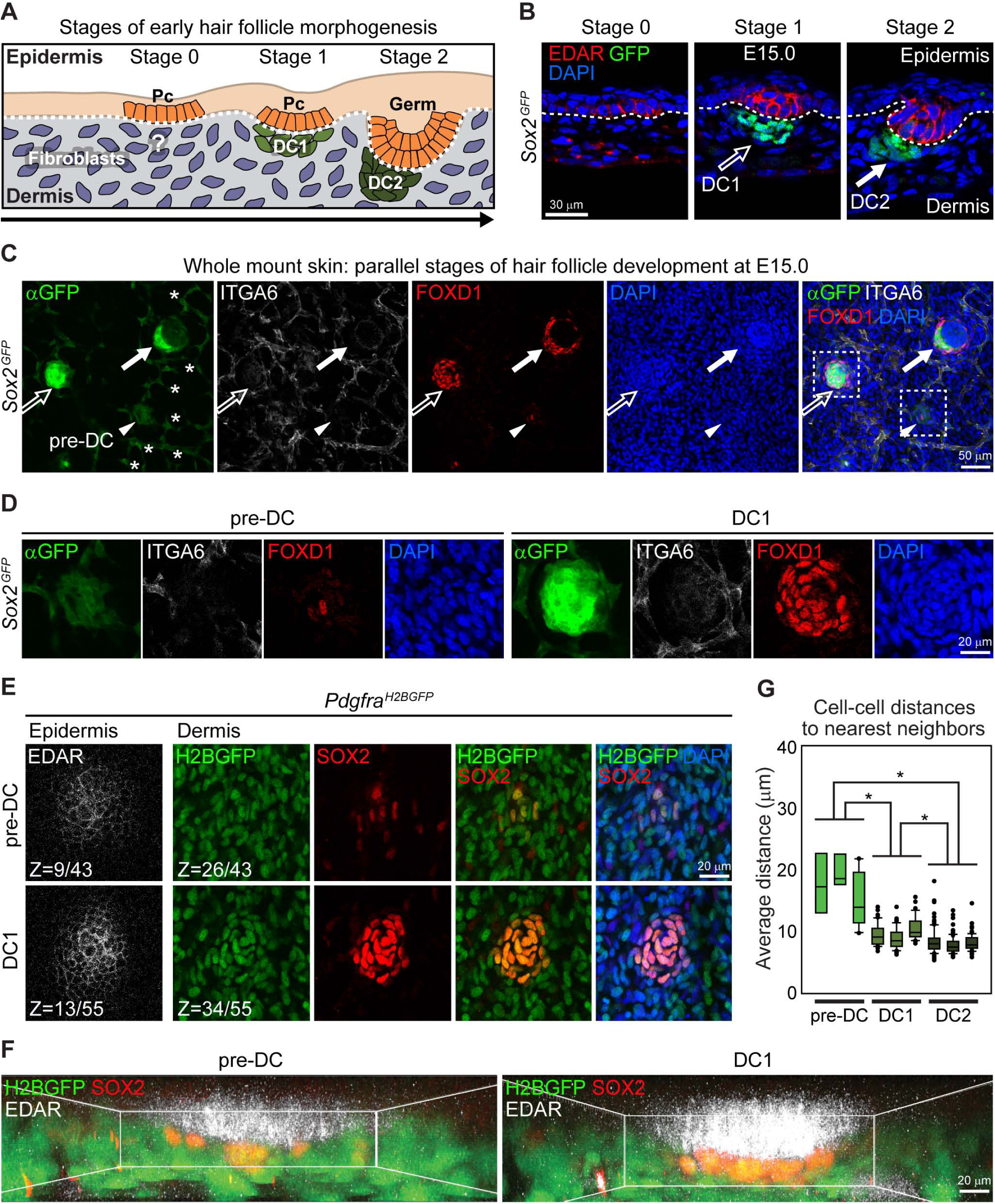
Identification of Unclustered Precursors of Dermal Condensates. (A) Schematic depicting early key stages of hair follicle morphogenesis. Initiation of hair follicle placodes (Pc) precedes formation of the clustered dermal condensate (DC) niche. Whether DC fate specification occurs before aggregation is unclear. (B) Immunofluorescence for placode marker EDAR on a sagittal section of E15.0 back skin from *Sox2*^*GFP*^ embryos. GFP high signal marks the DC of stage 1 (DC1, empty arrow) and stage 2 hair follicles (DC2, filled arrow). Note parallel hair follicle stages 0, 1 and 2 at E15.0. Dotted line demarcates basement membrane. Red speckles in dermis are non-specific. DAPI marks all nuclei. (C) Immunofluorescence whole mount staining for GFP (*Sox2*^*GFP*^ reporter), Schwann cell marker ITGA6 and DC marker FOXD1 on E15.0 *Sox2*^*GFP*^ back skin. Note parallel hair follicle development stages in the top view of a low magnification confocal scan. GFP high signal is in DC1 and DC2. DC2 shows already typical anterior-posterior polarized downgrowth. The GFP low signal marks a group of unclustered, FOXD1^+^ potential DC precursors (pre-DC, arrowhead). GFP low also marks the ITGA6^+^ Schwann cell network (asterisks). (D) Magnification of inset from (C) for pre-DC (arrowhead) and DC1 (open arrow). Pre-DC appears unclustered. DC1 shows typical arrangement of clustered nuclei. (E) Immunofluorescence whole mount staining for EDAR and SOX2 on E15.0 *Pdgfra*^*H2BGFP*^ back skin. Nuclear H2BGFP is in mesenchymal cells. Z-sections from a 3D-confocal scan are shown at the epidermal and dermal level. SOX2^+^ unclustered pre-DC and DC1 cells are underneath EDAR^+^ placodes and are fibroblast-type, expressing mesenchymal H2BGFP. Z values are section # of total optical sections in scan from epidermis into dermis. Section intervals = 1 μm. (F) 3D reconstructed images of pre-DC and DC1 generated by Imaris using the confocal scans in (E). (G) Quantification of cell density of pre-DC, DC1 and DC2. Average distances of nearest 5 neighboring nuclei for each pre-DC, DC1 and DC2 were measured in 3D reconstructions in (F) and Figure S1E. n = 3 for pre-DC, DC1 or DC2. Data are mean ± SD from two embryos. *p < 0.01.

The morphological detection of the DC after placode initiation (Chen et al., 2012), combined with discovery of DC markers (Botchkarev et al., 1999; Chen et al., 2012; Clavel et al., 2012; Huh et al., 2013) and the definition of a DC gene expression signature (Sennett et al., 2015) has led to the current model of niche fate specification in response to progenitor signal(s) that coincides with condensate formation. However, the identification of unclustered dermal cells with high and localized Wnt activity beneath presumptive placodes (Zhang et al., 2009), and low-level expression of DC signature genes in immunophenotypically non-DC dermal cells (Sennett et al., 2015) raise the possibility of DC niche fate acquisition prior to cluster formation. Contextualizing these early events in hair follicle niche formation is further complicated by the rapidity with which morphogenesis occurs; Wnt signaling activity appears virtually simultaneously in Pc and DC (Tsai et al., 2014; Zhang et al., 2008), rendering parsing out the precise order and timing difficult. Recent work also demonstrated direct mesenchymal self-organization ability in the absence of an epidermal pre-pattern (Glover et al., 2017) further calling into question the origin and timing of niche fate specification (**Figure 1A**): 1) is the niche fate specified before niche cluster formation and signaling function and 2) how is the timing of niche fate acquisition related to progenitor fate and signaling?

Here, by combining 3D and 4D time-lapse live imaging with fluorescence-activated cell sorting, bulk and single-cell transcriptomics, proliferation kinetics, and mutant mouse models, we identify fated precursors of the clustered DC niche prior to signaling center formation. Based on a molecular time-lapse of dynamically changing niche gene signatures and ordering the cell fates in developmental pseudotime we define the cell fate trajectory from fibroblasts through DC niche precursors that are fated to become the clustered niche. In this process, DC precursors (pre-DC) undergo a transitional molecular state that is consolidated towards the DC niche biological fate through waves of transcription factor and signaling molecule upregulation. Finally, with epidermal Wnt signaling ablation that prevents placode formation, and with deletion of placode-derived FGF20 that precludes DC aggregation we demonstrate dependence of niche cell precursor acquisition on placode progenitor fate.

## RESULTS

### Identification of Unclustered Precursors of Dermal Condensates

Previous studies have demonstrated cell aggregation is the cellular mechanism inciting DC formation (Biggs et al., 2018; Glover et al., 2017), but when and how the DC niche fate is acquired remains unknown. Here, to identify potential DC-fated precursors before cluster formation and niche establishment, we followed up on our previous observation of a mixed population of Schwann and DC-like cells isolated from embryonic *Sox2*^*GFP*^ reporter skin (**Figure S1A**) (Sennett et al., 2015), that expresses low levels of GFP along with a variety of DC and Schwann cell genes (**Figures S1A, S1B** and **S1C**). GFP (**Figures 1B** and **S1A**) and *Sox2* (**Figure S1B**) are highly expressed in early DC (Clavel et al., 2012; Driskell et al., 2009; Sennett et al., 2015), where condensed cells are visible underneath flat and barely invaginated EDAR^+^ hair placodes of stage 1 hair follicles (**Figure 1B**, open arrow) (Paus et al., 1999; Pispa et al., 2008; Sennett et al., 2015). GFP levels remain high after initial placode downgrowth and polarization in stage 2 hair follicles (**Figure 1B,** arrow). We named these two DC stages with high *Sox2* expression DC1 and DC2 accordingly, and surmised that cells with low *Sox2*^*GFP*^ expression could represent a precursor state. However, difficulty of detecting low GFP levels, even in Schwann cells, in sagittal skin sections precluded their convincing discovery (**Figure 1B**). To detect GFP low cells with high sensitivity we performed multicolor immunofluorescence in whole mount skin of E15.0 *Sox2*^*GFP*^ embryos (**Figure 1C)**. As predicted from sagittal sections, in the top view GFP was strong in DC1 and DC2, but with anti-GFP staining, low GFP levels were now detectable also in Schwann cells, highlighted by ITGA6, forming a GFP low/ITGA6^+^network within the developing skin (**Figure 1C,** asterisks). Importantly, we found few loosely arranged GFP low/ITGA6^--^cells in the dermis in a location fitting of a hair follicle pattern (**Figure 1C,** arrowhead). Co-labeling with the previously identified DC marker FOXD1 (**Figure S1C**) (Sennett et al., 2015) labeled DC1, DC2 and these GFP low cells, suggesting that they might be DC precursors (pre-DC) before DC niche formation. High magnification images suggested that these cells are still unclustered, unlike the clearly visible cell aggregations of DC1 stage hair follicles (**Figure 1D**).

To demonstrate that potential pre-DC cells are associated with hair follicles we next performed whole mount immunofluorescence for SOX2 and EDAR on E15.0 *Pdgfra*^*H2BGFP*^ back skin. *Pdgfra* is highly expressed in fibroblast-type mesenchymal cells (**Figure S1C**) (Collins et al., 2011; Hamilton et al., 2003), which are marked in this reporter line by nuclear GFP. In optical slices of 3D confocal scans, SOX2^+^ pre-DC cells at the dermal level were underneath EDAR^+^ placodes at the epidermal level and appeared to be unclustered compared to aggregated DC1 and DC2 (**Figures 1E** and **S1D**). Overlap of SOX2 with nuclear H2BGFP demonstrated the fibroblast-type of potential pre-DC cells. 3D reconstructions further illustrated the unclustered state of pre-DC cells **(Figure S1E, Supplemental Movies 1-3)**. Z-projections demonstrated the close proximity of unclustered pre-DC to the overlying placode (**Figure 1F**). Finally, to confirm that pre-DC cells are truly unclustered we measured, in 3D, the distance of each nucleus to its five nearest neighbors for pre-DC, DC1 and DC2 (**Figure 1G**). Cells in DC2 clusters were significantly closer to each other compared to all others. Pre-DC cells, however, were significantly further apart compared to those of DC1 and DC2, confirming their loose initial arrangement. As other H2BGFP^+^ fibroblasts were interspersed between pre-DC cells (**Figure 1E**), it suggests that early fate acquisition does not have to be synchronized among direct neighbors. In summary, our data thus far identified the existence of uncondensed potential DC precursor cells suggesting that the DC fate is specified prior to DC formation and function.

### DC Precursor Cells Are in a Transitional Transcriptional State Towards a DC Niche Fate

After identifying potential DC precursors in stage 0 hair follicles with SOX2 protein, GFP reporter and the DC signature gene FOXD1, we next sought to isolate this population for exploring the molecular controls of DC fate acquisition and its niche functions. The simultaneous presence of pre-DC, DC1 and DC2 from stage 0, stage 1 and 2 hair follicles, respectively, in E15.0 back skin provided us with an excellent opportunity to compare multiple parallel developmental stages in isolated cells. For this, we modified our previous DC isolation method (**Figure S1A**) (Sennett et al., 2015) to separately purify *Sox2*^*GFP*^ high DC1 and DC2 by differential DC marker CXCR4 expression (Sennett et al., 2014) in DC2 (arrow) compared to DC1 (open arrow) (**Figure 2A** and **2B**). Simultaneously we stained for mesenchymal marker PDGFRA to isolate dermal fibroblasts (Fb), for ITGA6 to obtain Schwann (Sch) cells and separately isolate pre-DC as the *Sox2*^*GFP*^ low/PDGFRA^+^/CXCR4^-^population. Additionally, PDGFRA^-^/Sox2^GFP-^cells were sorted as a negative (Neg) population containing all other remaining skin cells (**Figure 2B**). To confirm enrichment of Fb and DC we performed qRT-PCR of known marker genes (**Figure 2C**). As expected, *Lum* (Lumican) was highly enriched in fibroblasts, while *Cxcr4* was absent in pre-DC and detected at increasing levels in DC1 and DC2, and *Sox2* expression was low in both pre-DC and Schwann cells and high in DC1 and DC2. Importantly, Schwann cell marker *Fabp7* showed strong enrichment in Schwann cells and not in isolated pre-DC (**Figure S2A**), and all other Fb/DC markers were also absent (**Figure 2C**). Finally, *Foxd1* was highly expressed at all three DC stages, and not in Fb or Schwann cells, confirming successful isolation of pre-DC cells in agreement with our 3D imaging analysis.

**Figure 2.**
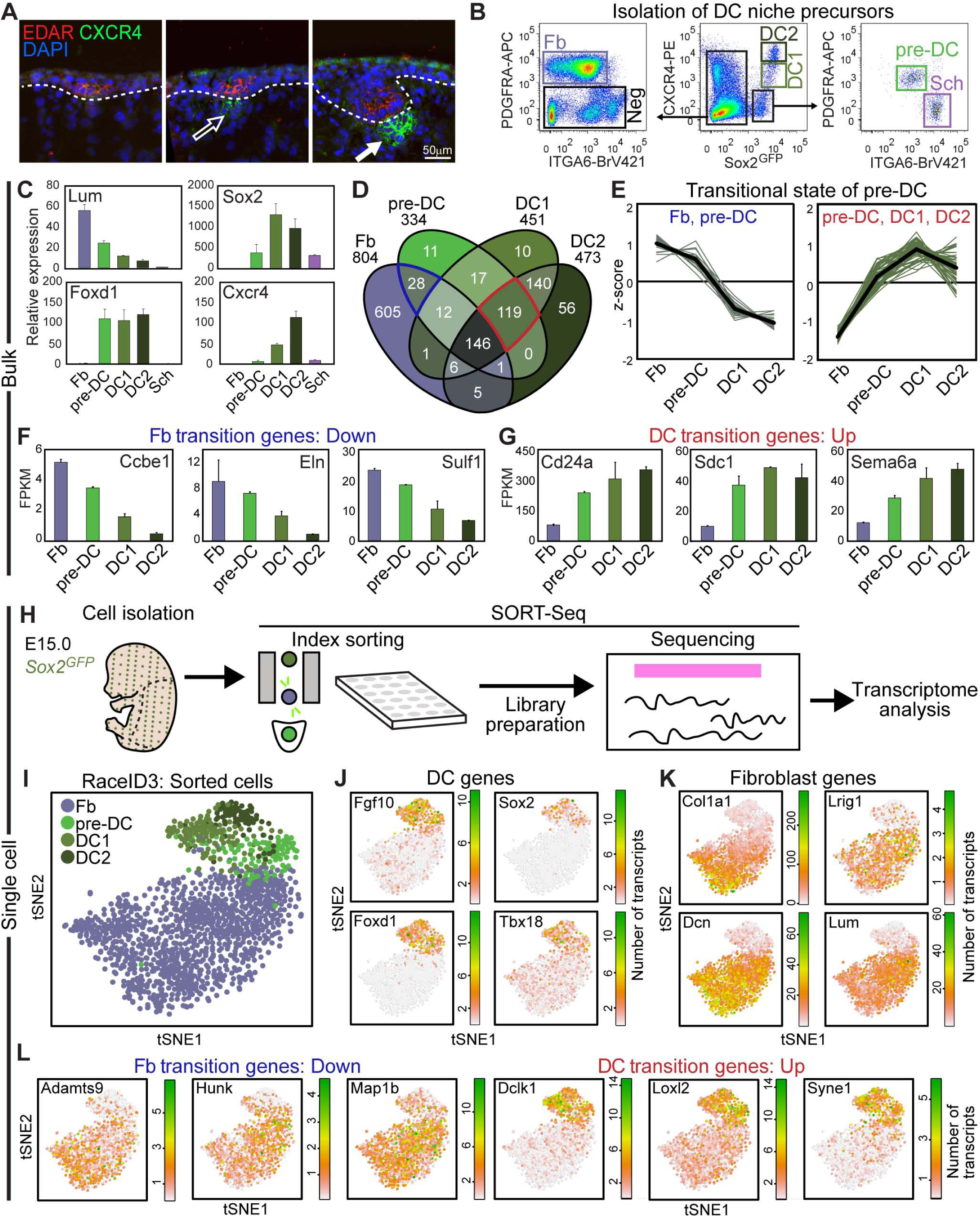
DC Precursor Cells Are in a Transitional Transcriptional State Towards a DC Niche Fate. (A) Immunofluorescence for DC marker CXCR4 and placode marker EDAR in E15.0 back skin. CXCR4 is expressed at higher levels in DC2 (filled arrow) compared to DC1 (open arrow). Dotted line marks basement membrane and DAPI marks all nuclei. (B) FACS isolation of pre-DC, DC1, DC2, as well as fibroblasts (Fb), Schwann cells (Sch), and a population of negative cells (Neg) from live, EpCAM^-^ cells. (C) qRT-PCR verification of marker genes in isolated Fb, pre-DC, DC1, DC2 and Sch cells. (D) Venn diagram of gene signatures in fibroblasts, pre-DC and two DC stages. All genes are enriched compared to Sch and Neg. The overlaps represent commonly enriched genes in corresponding populations compared to all others. Blue outlines increased overlap between Fb/pre-DC, compared to Fb/DC1 or Fb/DC2. Red outlines large overlap of pre-DC with mature DC stages. (E) Transitional transcriptional state of pre-DC. Left: Z-score normalized expression of Fb/pre-DC overlap genes (blue) in Fb, pre-DC, DC1, and DC2 reveal progressive downregulation of Fb genes. Right: Z-score normalized expression of overlap of pre-DC/DC1/DC2 (red) and 58 established DC signature genes shows increased DC gene expression (Sennett et al., 2015). Pre-DC has intermediate levels for both Fb and DC genes. (F and G) FPKM barplots of representative transitional genes. (H) Experimental strategy for combined single cell sorting and single cell transcriptome analysis by SORT-seq. Fb, pre-DC, DC1 and DC2 were FACS-purified as in (A). Each index sorted cell is connected to its transcriptome through the plate position. (I) tSNE map of all sorted cells color-coded by cell type. (J) Expression of representative DC signature genes projected onto the tSNE map from (I). Transcript counts are in linear scale. (K) Expression of representative Fb signature genes projected onto the tSNE map from (I). (L) Expression of representative transitional Fb and DC genes from the Fb/pre-DC and pre-DC/DC1/DC2 overlaps projected onto the tSNE map from (I).

We then proceeded with population-based transcriptome analysis that provided high sensitivity of gene signature discovery, as previously described (Rezza et al., 2016; Sennett et al., 2015). Bulk RNA-sequencing (RNA-seq) was performed with low RNA amounts obtained from all isolated populations. We first globally compared the transcriptomes by principal component analysis of the top variably expressed genes (**Figure S2B**) and hierarchical clustering (**Figure S2C**), confirming the similarities between DC1 and DC2 and establishing multiple co-regulated gene groups within all six populations. Interestingly, pre-DC was positioned between Fb and DC1 suggesting pre-DC dually share characteristics of both dermal fibroblasts and clustered DC. We next calculated significantly differentially expressed genes (DEGs) in each population by ANOVA (false discovery rate < 0.05) and compared the resulting total of 6,454 genes. We then established gene signatures using significant DEGs and selecting for genes with FPKM ≥ 1 and an expression fold change ≥ 2, which resulted in 1,890 signature genes for all six cell types. To focus on the potential pre-DC cells and their relationship to fibroblasts and the condensate niche, we analyzed gene expression relationships in a four-way Venn diagram analysis all significantly increased genes in Fb, pre-DC, DC1 and DC2 compared to Sch and Neg populations (**Figure 2D** and Table S1). A total of 1,157 genes were enriched in the four mesenchymal populations, including 692 population-unique signature genes. However, we also found several interesting population overlaps. As the DP and its embryonic DC predecessor are fibroblast-derived mesenchymal cells (Driskell et al., 2013; Jinno et al., 2010), it was revealing to note that the unique pre-DC overlap with Fb (28 genes) was larger than both the unique overlaps of Fb and DC1 (1 gene) and Fb and DC2 (5 genes) (**Figure 2D**, blue), suggesting that pre-DC shares transcriptional similarity with fibroblasts fitting with a Fb-to-DC trajectory. Indeed, exploring the expression trajectory of genes shared between Fb and pre-DC demonstrated a transition of high-to-low expression from Fb to DC with intermediate levels in pre-DC (**Figure 2E**). Several genes included extracellular matrix (ECM)-components such as *Ccbe1* and *Eln*, and ECM-modifiers such as *Sulf1* (**Figure 2F**). Conversely, differential expression analysis confirmed a high overlap of 119 genes between pre-DC, DC1, and DC2 (**Figure 2D,** red), which included 58 previously described DC signature genes (Sennett et al., 2015). Also here, pre-DC expressed many of these shared DC signature genes at intermediate levels between fibroblasts and DC1/DC2, suggestive of its developmental precursor status to morphologically identifiable DC (**Figures 2E** and **2G**).

We next complemented population-based bulk RNA-sequencing with single cell transcriptome analyses that allow for exploration of minute cell-to-cell transcriptional changes that define cell heterogeneity and differentiation (Grün et al., 2015; Haber et al., 2017; Joost et al., 2016; Yang et al., 2017). Importantly, it also enables a high-resolution definition of lineage relationships along a developmental trajectory (Artegiani et al., 2017; Herman et al., 2018), with which we can corroborate the developmental link of pre-DC between fibroblasts and DC1/DC2. For this, we used Sorting and Robot-Assisted Transcriptome Sequencing (SORT-Seq) (Muraro et al., 2016) with our established DC cell isolation strategy. SORT-seq is a powerful approach combining live cell isolation of identified, known single cells with their transcriptome information through an index, connecting sorted cells with single cell RNA-seq wells (**Figure 2H**). We isolated, index sorted, and sequenced 1,371 fibroblasts, 149 pre-DC, 152 DC1, and 151 DC2. Of those, 1,076 fibroblasts, 107 pre-DC, 118 DC1, and 82 DC2 were used for analysis based on a minimum requirement of 6,000 transcripts/cell and established quality controls to exclude poorly sequenced cells. Based on population frequencies observed by flow cytometry it would require ∼15,000 total unenriched cells to obtain similar transcriptome information of our sorted pre-DC cells (150x enrichment). We then used the Rare Cell Type Identification 3 (RaceID3) clustering algorithm that was previously used in various iterations to identify rare intestinal cell types and lineage relationships between stem cells and differentiated cells (Grün et al., 2015, 2016; Herman et al., 2018). RaceID3 uses k-medoid clustering of the transcriptome-based correlation matrix that is visualized in two dimensions using t-distributed stochastic neighbor embedding (tSNE) (Van Der Maaten and Hinton, 2008) to show cell-cell gene expression similarities. Superimposing the Fb and DC cell identities onto the RaceID3-generated tSNE revealed clear distinctions between all isolated populations confirming our cell isolation strategy (**Figure 2I**). At the same time, it highlighted similarities between all DC stages and placed pre-DC between fibroblasts and DC1/DC2, similarly suggesting a lineage trajectory from Fb to DC through an intermediate pre-DC fate transition. As expected, all sorted DC populations expressed established DC signature genes, including *Fgf10* (Mailleux et al., 2002; Suzuki et al., 2000), *Sox2, Foxd1,* and *Tbx18* (Grisanti et al., 2013) (**Figure 2J**), and sorted fibroblasts expressed established fibroblast markers, including *Col1a1, Lrig1* (Driskell et al., 2013), *Dcn* (Ferdous et al., 2010; Philippeos et al., 2018), and *Lum* (Philippeos et al., 2018) (**Figure 2**K). Also here, cursory exploration of expression patterns revealed gradual downregulation of Fb genes and upregulation of DC genes (**Figure 2L**). Altogether, these bulk and single cell transcriptome data from purified embryonic skin cells indicate that the unclustered pre-DC precursors are at a transcriptionally transitional state from fibroblasts towards a permanent DC niche fate.

### DC Precursors are Developmental Intermediates Between Fibroblasts and The Mature DC Niche

As the identified pre-DC cells express genes characteristic of both fibroblasts and DC, we hypothesized that they are precursors of the mature DC niche at an intermediate developmental state between fibroblasts and DC. To explore this lineage trajectory, we first utilized RaceID3 to identify distinct clusters of single cells with similar transcriptome as a basis for calculating lineage relationships between related cell types, and visualized them in t-SNE (**Figure 3A**). Clusters 6, 7 and 8 were highly similar in shape and distribution to our sorted pre-DC, DC1 and DC2 cells, respectively (**Figures 2H** and **3A**), again validating our sorting strategy. Calculation of the overlap percentages confirmed that clusters 6, 7, and 8 were mostly comprised of pre-DC, DC1 and DC2, respectively (**Figure S3A**). We then applied the Stem Cell Identification (StemID2) algorithm (Grün et al., 2016; Herman et al., 2018) to order the RaceID3-identified clusters by populating the links between the clusters’ medoids with cells from the linked clusters based on degree of transcriptional similarity (**Figure 3B**). StemID2 enables postulating differentiation trajectories and stem cell identity from single cell RNA-sequencing data (Grün et al., 2016; Herman et al., 2018). Amongst DC clusters (clusters 6, 7, 8), the pre-DC-enriched cluster 6 was closer to the fibroblast clusters. Amongst the sorted fibroblasts, fibroblast cluster 5 was the sole calculated link to pre-DC cluster 6, suggesting that pre-DC cells emerged from those closely related fibroblasts (**Figure 3B**). It also demonstrated that pre-DC are the link for transition into DC1/DC2, which were connected by StemID2 to pre-DC with equal confidence. To independently confirm this developmental trajectory, we turned to an alternative approach, using the Scanpy toolkit (Wolf et al., 2018) for Louvain clustering and diffusion maps. Louvain clustering, a method to extract communities from large networks (Blondel et al., 2008), with the same 1,383 cells as used for RaceID3 clustering, also showed clear distinctions between all sorted populations, placing again pre-DC between fibroblasts and DC1 (**Figures 3C** and **S3B**). Importantly, applying diffusion pseudotime, a method for ordering cells along lineage trajectories toward different cell fates (Haghverdi et al., 2016), clearly demonstrated the lineage path of niche specification from nearest fibroblasts (Louvain cluster 5) through pre-DC towards DC1 and DC2 (**Figures 3D, S3C**).

**Figure 3.**
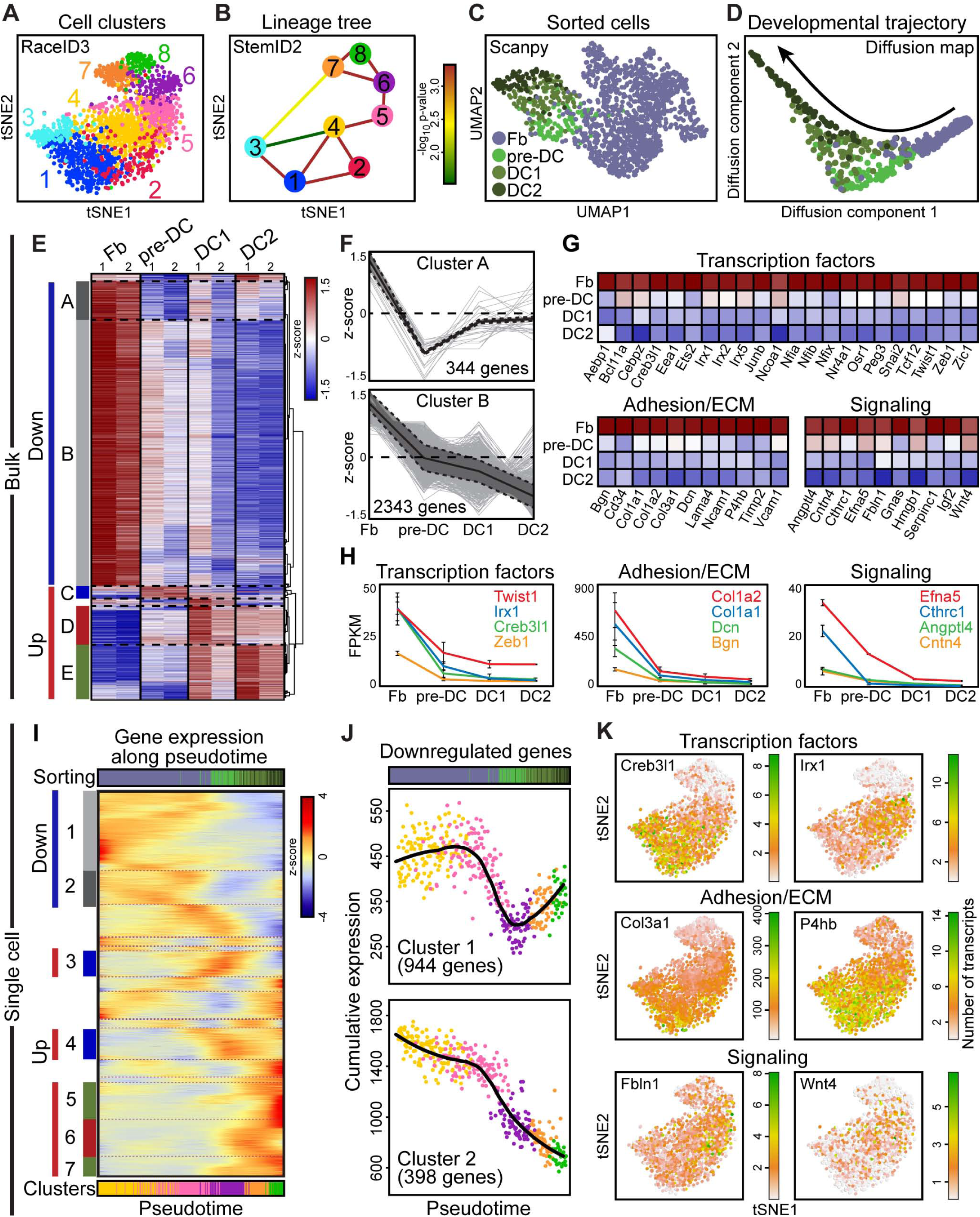
DC Precursors are Developmental Intermediates Between Fibroblasts and The Mature DC Niche. (A) tSNE map from Figure 2I color-coded according to RaceID3 k-medoids clustering. (B) StemID2 lineage tree of RaceID3 k-medoids clustering output. Color of the line connecting clusters indicates p-value of link. (C) UMAP plot of Louvain clustering (Scanpy) of Fb, pre-DC, DC1, DC2. Sorted cell identities are color-coded by cell type. (D) Diffusion map highlighting the developmental trajectory from Fb through pre-DC to DC2 and color-coded by cell type. (E) Trend analysis by Spearman rank clustering of differentially expressed genes from Fb to pre-DC, DC1 and DC2 in bulk RNA-seq. (F) Expression trajectories of downregulated genes from Fb to pre-DC. Solid black lines are average z-score flanked by dotted black lines representing SD. (G) Heatmaps of relative expression of downregulated transcription factors, adhesion/extracellular matrix, and signaling genes across the DC developmental trajectory. (H) Absolute FPKM expression levels of representative genes from each category in (E). (I) Self organizing map of co-regulated gene clusters across the plotted pseudotime of ordered individual cells based on the gradual transition of their transcriptomes. Bottom bar represents ordered cells color-coded by RaceID3 cluster in (A). Top bar highlights ordered cells based on sorted identities in (C). (J) Cumulative expression plots of downregulated genes along pseudotime. Each dot represents the cumulative expression of all genes per cell in a given cluster from (I). The black line is the running average across pseudotime. (K) Expression of representative transcription factors, adhesion/ECM molecules, and signaling factors from gene clusters 1 and 2 projected onto the tSNE map from Figure 2I.

Having established by single cell RNA-sequencing analyses the differentiation trajectory from Fb to DC2, we next re-examined the bulk RNA-sequencing for gene expression trends. For this we performed hierarchical clustering on all genes that had a >2 fold difference between maximum and minimum FPKM of any 2 populations. We noted four major trends: genes with highest expression in fibroblasts (down A and B), and genes with highest expression in pre-DC (up C), DC1 (up D) and DC2 (up E) (**Figure 3E** and Table S2). 2,687 genes were downregulated from fibroblasts through DC2 (**Figure 3F**). Closer examination of these genes revealed major downregulation of many Fb transcription factors, adhesion/ECM molecules, and signaling factors (**Figures 3G** and **3H**) suggesting their involvement in the rapid and sustained loss of fibroblast function as the DC fate is acquired and throughout DC differentiation.

Finally, we utilized our single cell transcriptome data to precisely define the pseudotemporal ordering of fibroblasts and DC populations along the differentiation trajectory and explore patterns of gene regulation during niche fate acquisition. For this, we devised a self-organizing map using the fibroblast clusters closest to DC and the DC-enriched clusters (**Figure 3I**). We excluded *Dlk1* expressing fibroblast clusters 1-3 that likely represented reticular fibroblasts of the lower dermis unrelated to DC and hair formation (**Figure S3D**) (Driskell et al., 2013), and included fibroblast clusters 4 and 5 as both were intermingled in Louvain cluster 5 and closest to pre-DC-enriched cluster 6 (**Figure S3E**). Pseudotime gene expression analysis revealed cognate but more refined trends to those observed in the bulk RNA-sequencing (**Figure 3I** and Table S3): genes with their highest expression in fibroblasts (down clusters 1, 2), and genes with their highest expression in pre-DC (up clusters 3, 4), DC1 (up cluster 6), and DC2 (up clusters 5, 7). Amongst the 2,687 genes downregulated from fibroblasts through DC2 in population-based RNA-sequencing (**Figure 3F**) and the 1,342 genes identified in the single cell transcriptomes (**Figure 3J**), 951 genes were in common (35% of bulk-identified targets, 70% of single cell-identified targets). Also here, these included several transcription factors, adhesion and ECM components, and signaling molecules necessary for fibroblast function (**Figure 3K**). *Creb3l1* is a secreted factor essential for collagen metabolism (Keller et al., 2018), while *Irx1* and *Twist1* are transcription factors that control expression of matrix-stiffening components (García-Palmero et al., 2016), and *Zeb1* is known for promoting expression of vimentin (Liu et al., 2008). Additionally, expression of numerous collagens such as *Col1a1, Col1a2*, and *Col3a1*, as well as other ECM components such as *Dcn* and *Bgn*, and *P4hb*, an enzyme essential for procollagen synthesis (Winter and Page, 2000), was decreased between fibroblasts and DC populations. Finally, we identified signaling molecules that are uniquely expressed in fibroblasts, such as *Efna5, Cthrc1, Angptl4, Cntn4, Fbln1,* and Wnt4 that are all downregulated on the path to DC lineage differentiation. Altogether these data suggest that fibroblasts quickly halt transcription of fibroblast-associated genes as they transition through the pre-DC intermediate into the definitive DC niche fate.

### Loss of Proliferation is Concomitant with Acquisition of DC Fate

It has been well established that niche cells in mature dermal papillae are quiescent and very rarely divide (Moffat, 1968; Tobin et al., 2003). Also the aggregated DC, as the embryonic precursor of the DP (Grisanti et al., 2013), was recently shown to be largely non-proliferative (Biggs et al., 2018). Our discovery of unclustered pre-DC cells then provided an opportunity to fine-map the timing of cell-cycle exit with respect to niche fate acquisition. As predicted, many genes involved in the molecular control of cell proliferation, including major cell cycle components, were downregulated from fibroblasts to DC2 (**Figure 4A**). Interestingly, gene set enrichment analysis revealed a significant correlation of proliferation-associated genes in fibroblasts compared to pre-DC, suggesting that the exit of proliferation could be as early as during pre-DC fate acquisition (**Figure 4B**). Indeed, in pseudotime-ordered cells a rapid declining expression of pro-proliferative genes such as *Rbbp7, Mki67*, and *Pcna* coincided with the sharp upregulation of cell-cycle inhibitors, such as *Cdkn1a* and *Btg1* (Zhu et al., 2013). Importantly, both events occurred prior to the emergence of pre-DC and remained sustained through DC2 (**Figures 4C** and **4D**), suggesting that cell cycle exit is a feature of pre-DC fate acquisition.

**Figure 4.**
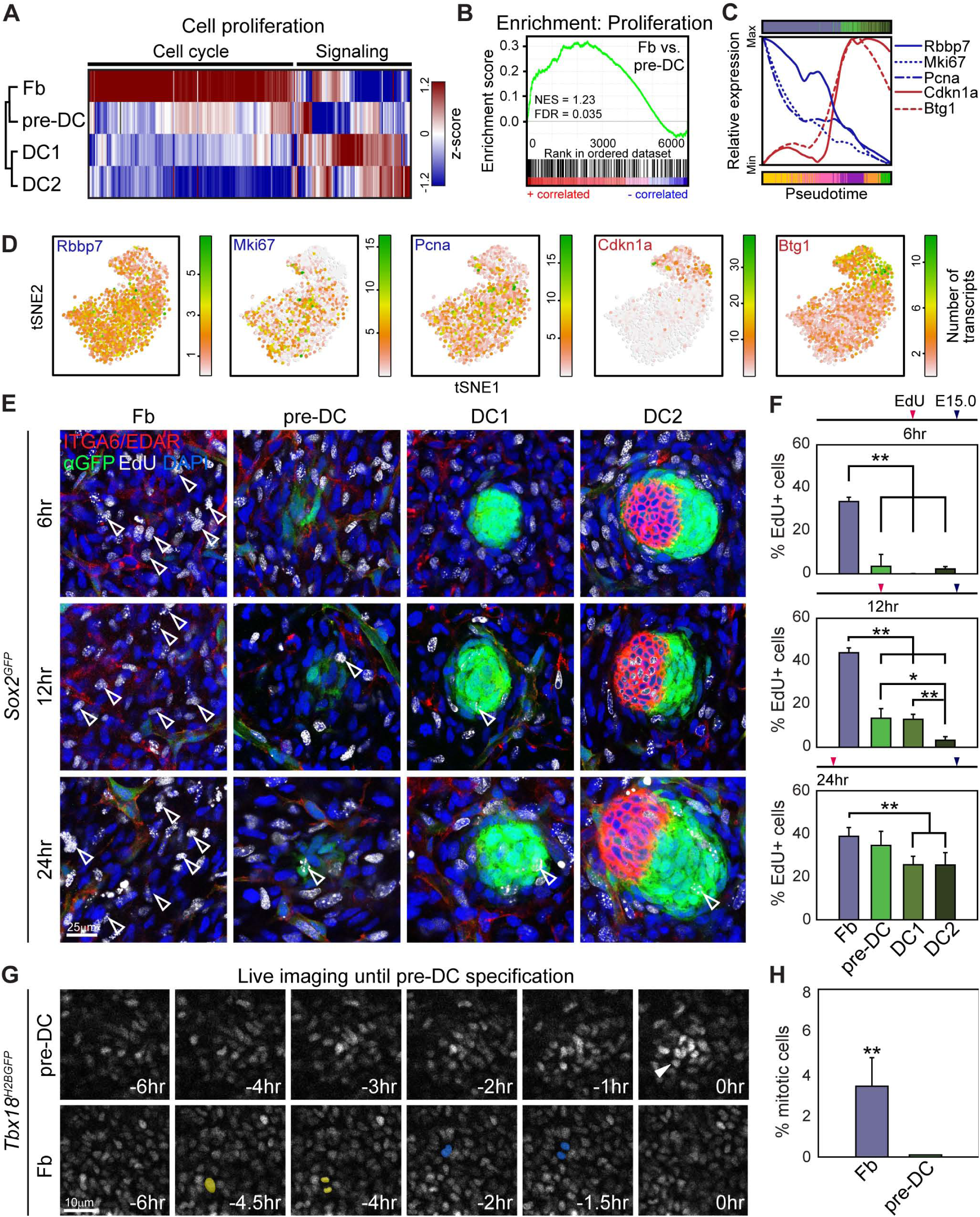
Loss of Proliferation is Concomitant with Acquisition of DC Fate. (A) Hierarchical clustering of cell proliferation gene expression in bulk-isolated Fb, pre-DC, DC1, and DC2. Cell cycle genes are shut down from Fb to pre-DC through DC2. (B) Gene Set Enrichment Analysis for cell proliferation genes shows positive correlation for Fb and negative correlation for pre-DC. (C) Normalized running means of expressed transcript numbers in pseudotime ordered cells for pro-proliferative (*Rbbp7, Mki67, Pcna*) and anti-proliferative (*Cdkn1a, Btg1*) genes. (D) Expression of pro-proliferative and anti-proliferative genes projected onto the tSNE map from Figure 2I. (E) EdU pulse-chase experiment for proliferation rates in Fb, pre-DC, DC1 and DC2. EdU was injected 6h, 12h, or 24h prior to harvest and analysis of E15.0 *Sox2*^*GFP*^ skin by immunofluorescence for GFP, ITGA6 and EDAR. Representative z-sections from confocal scans are shown for interfollicular Fb and pre-DC, DC1, DC2. Examples of EdU^+^ proliferating cells are highlighted by empty arrowheads. (F) Quantification of EdU^+^ Fb, pre-DC, DC1 and DC2. Note drastic drop of proliferation rate at 6h time point from Fb to DC precursors, sustained through DC2. n = 3 interfollicular Fb areas, pre-DC, DC1 and DC2. Data are mean ± SD from two embryos. *p < 0.05, **p < 0.01. (G) Representative stills from time-lapse live imaging in *Tbx18*^*H2BGFP*^ embryonic skin for detection of cell divisions. Pre-DC cells were identified as H2BGFP high cells after transitioning from fibroblasts and backtracked in time for 6 hours. Fibroblasts were H2BGFP low. Note absence of cell divisions in pre-DC through 6 hours before fate acquisition. Two examples for mitotic cell divisions in Fb are pseudo-colored in yellow or blue. (H) Quantification of mitotic cells during 6 hour tracking. n = 2. Data are mean ± SD from two embryos. **p < 0.01.

To independently test when proliferation exit occurs relative to DC fate acquisition, we performed EdU proliferation assays to mark proliferating cells during the S-phase of the cell cycle (**Figures 4E** and **S4A**). EdU injection 6 hours prior to E15.0 analysis showed that 33% of fibroblasts were proliferating (**Figure 4F**). By contrast, less than 3% of DC1 and DC2 were going through S-phase suggesting that the vast majority of maturing DC cells had exited the cell cycle. Notably, only 3% of pre-DC precursors incorporated EdU suggesting that already during this early step of DC fate acquisition cells shut down proliferation. Analyzing 12 and 24 hour chases showed increased EdU incorporation in pre-DC and DC1 fated cells, likely reflecting their developmental history coming from actively proliferating fibroblasts at the time of EdU injection. Interestingly, with the 12 hour EdU chase the percentage of proliferating cells in DC2 was significantly lower than in pre-DC and DC1 and as low as 6h levels. Given that DC1 is progressing to DC2 during this 12 hour chase, the results herein suggest that progression to DC2 from DC1 does not require cell proliferation.

Finally, to confirm that exit of proliferation is already occurring during DC niche fate establishment before pre-DC precursor detection, we tracked with 4D time-lapse live imaging the cellular processes up to pre-DC fate specification (**Figure 4G**). For this we utilized live imaging of embryonic ex vivo back skins from *Tbx18*^*H2BGFP*^ mice that express high levels of nuclear GFP in the DC (Grisanti et al., 2013). Image analysis of DC1 and DC2 confirmed high GFP expression as expected (**Figure S4B**). Importantly, pre-DC cells also already express high nuclear GFP, while fibroblasts exhibit low GFP expression levels (**Figure 4G**). Tracking cell divisions during a 6 hour period prior to pre-DC fate acquisition detected actively-proliferating interfollicular fibroblasts as predicted (**Figure 4G** and **4H**). By contrast, no pre-DC cells were dividing during the same time span, indicating that proliferation exit is occurring during DC fate acquisition in the fibroblasts-to-pre-DC transition.

### Three Distinct Waves of Transcription Factor and Signaling Molecule Upregulation Occur Along the DC Niche Developmental Trajectory

Besides shutdown of proliferation and gradual loss of the fibroblast gene expression signature, a gradual upregulation of the DC niche signature occurs during DC niche fate acquisition (**Figures 2E, 2G, 2L**, 3E and **3I**). We next explored the precise timing of the molecular fate acquisition and consolidation from fibroblasts to pre-DC, through DC1 and DC2. We discovered three distinct waves of DC gene upregulation for pre-DC cell fate acquisition, DC1 clustering, and hair germ downgrowth stages at DC2, in both bulk (**Figure 5A**) and single cell (**Figure 5C** and Table S3) transcriptional analyses. The first wave was evident in gene clusters whose expression peaked early at pre-DC and then dropped, which likely includes factors necessary for fate acquisition and other early functions. Two additional waves of co-regulated gene clusters had sustained expression through DC2, but differed only in when their expression peaked: mid genes reached maximal expression in DC1, and late genes reached maximal expression in DC2. All three waves contained many transcription factors that could represent potential master regulators for DC niche specification and development (**Figure 5B-5G** and Table S4). Transcription factors expressed in the early wave included known key fate regulators such as *Twist2* (*Dermo1*) (Li et al., 1995), the Wnt signaling transcription factor *Lef1,* and the TGF-β effector Smad3 (**Figures 5B, 5D, 5F** and **S5A**). *Twist2* knock-out animals have sparse hair and thin skin, among many other abnormalities (Šošić et al., 2003). Similarly, *Lef1* knock-out results in abrogation of whiskers and back skin hair follicles, and of other ectodermal appendages such as mammary glands and teeth (van Genderen et al., 1994). We confirmed Twist2 expression in pre-DC cells by *in-situ* hybridization, which is shut down at the DC1 stage (**Figure 5D**). Many more transcription factors were in the mid gene cluster (**Figures 5E, 5F** and **S5B**) Mid-expressing transcription factors include *Trps1, Tshz1, Foxd1, Prdm1,* and *Sox18.* Mutations in *Trps1* are associated with Trichorhinophalangeal syndrome, a disorder characterized by hair follicle defects among others (Giedion et al., 1973), and Ambras Syndrome, a hypertrichosis disorder (Fantauzzo et al., 2008). *Prdm1* is a well-characterized marker for early papillary dermal development (Driskell et al., 2013; Horsley et al., 2006; Robertson et al., 2007). Mutations in *Sox18* have been shown to impact differentiation of adult dermal papillae (Villani et al., 2017). Finally, we resolved many transcription factors whose expression peaked in DC2, including Notch target genes *Hey1* and *Hes5*, as well as *Glis2, Foxp1*, and *Alx4* (**Figures 5E**, **5F** and **S5C**). *Foxp1* has noted functions in maintaining the quiescence of adult hair follicle stem cells (Leishman et al., 2013). Human mutations in *Alx4* have been associated with noted defects in hair follicle differentiation (Kayserili et al., 2009). In total, we discovered 11 early-wave, 16 mid-wave and 24 late-wave transcription factors (**Figure 5G**) that followed these temporally-restricted upregulation patterns along pseudotime, suggesting that a coordinated array of molecular changes is required for pre-DC to mature through DC1 and DC2.

**Figure 5.**
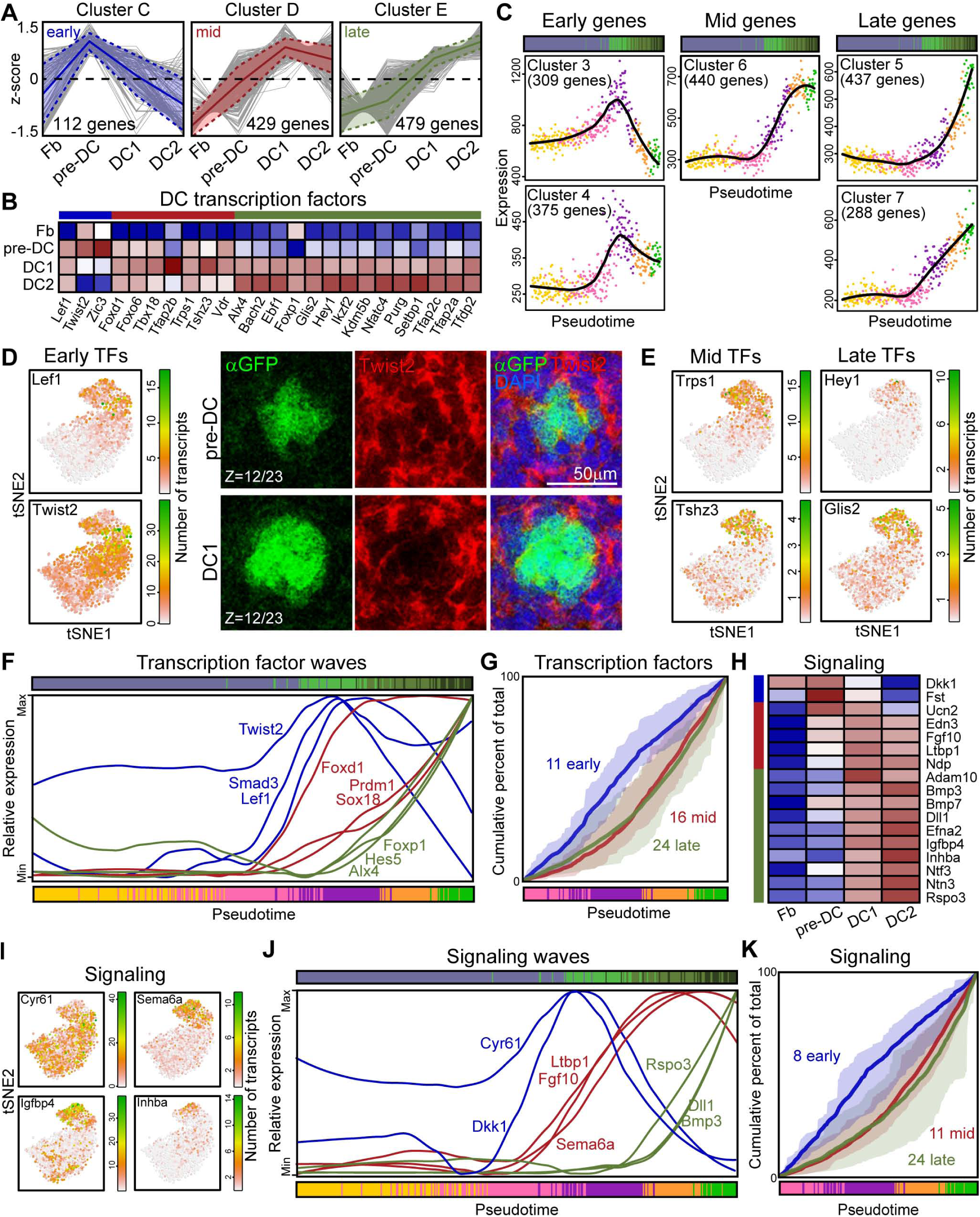
Three Distinct Waves of Transcription Factor and Signaling Molecule Upregulation Occur Along the DC Niche Developmental Trajectory. (A) Bulk RNA-seq expression trajectories of up-regulated genes from Fb to DC2 comprising Spearman rank clustered groups C (early, pre-DC peak), D (mid, DC1 plateau), and E (late, max at DC2) from Figure 3E. (B) Heatmap of relative expression (z-score) of transcription factors in groups C, D, and E (blue, red, green bars) from Figure 3E. (C) Cumulative expression plots of early, mid and late upregulated genes along pseudotime from Figure 3I. Each dot represents the cumulative expression of all genes per cell in a given cluster. The black line is the running average across pseudotime. (D) Left panel: Expression of early upregulated transcription factors projected onto the tSNE map from Figure 2I. Right panel: Parallel *in situ* hybridization for Twist2 and immunofluorescence for anti-GFP staining. Note Twist2 expression in pre-DC, which is downregulated in DC1. DAPI marks all nuclei. (E) Expression of mid and late upregulated transcription factors projected onto the tSNE map from Figure 2I. (F) Normalized running means of expressed transcript numbers in pseudotime ordered cells. Transcription factors are upregulated in three waves early during specification of DC precursors and consolidation of DC niche fate. (G) Cumulative percent of sum of upregulated expression of 51 transcription factors in early, mid, and late waves. Means and range of cumulative percent in each transcription factor wave are represented by thin solid and thick faded lines, respectively. (H) Heatmap of relative expression (z-score) of signaling-related genes in groups C, D, and E (blue, red, green bars). (I) Expression of representative upregulated signaling molecules projected onto the tSNE map from Figure 2I. (J) Normalized running means of expressed transcript numbers in pseudotime ordered cells. Signaling molecules are upregulated in three waves early during specification of DC precursors and consolidation of DC niche fate. (K) Cumulative percent of sum of upregulated expression of 43 signaling molecules in early, mid, and late waves. Means and range of cumulative percent in each transcription factor wave are represented by thin solid and thick faded lines, respectively.

As the dermal papilla acts as a major signaling niche for epithelial progenitor cells during postnatal hair growth and for stem cells in cycling adult hair follicles, we finally explored the expression and upregulation of signaling molecules during DC fate specification and niche formation. Also here, with our bulk and single cell RNA sequencing analyses we discovered a distinct pattern of signaling waves (**Figures 5H-5K** and Table S4). Signaling factors expressed in the short window of time around pre-DC include the Wnt inhibitor *Dkk1, Cyr61*, and *Fst* (**Figures 5I** and **5J** and **S5D**). *Cyr61*, known to induce senescence in fibroblasts, is also dynamically expressed in wounding healing to reduce fibrosis, likely through cell-cell adhesion interactions with integrins and heparin sulfate proteoglycans (Jun and Lau, 2010). *Fst* knock-out is associated with abnormal whisker growth (Jhaveri et al., 1998) and abrogated hair follicle development (Nakamura et al., 2003). The mid-acting signaling molecules include well known regulators *Fgf10*, the TGF-β modulator *Ltbp1*, and *Sema6a* (**Figures 5I** and **5J** and **S5E**); late-acting signaling molecules include key ligands such as ***Igfbp4, Inhba, Rspo3, Dll1,*** and *Bmp3* (**Figure 5I** and **5J** and **S5F**) – many of which continue to be expressed in the adult dermal papilla (Rezza et al., 2016). The global display as cumulative percent of total corroborates a resolution of 8 early, 11 mid, and 24 late upregulated signaling factors (**Figure 5K**). Overall these data suggest that development of DC, from fate acquisition in a subset of fibroblasts through aggregated DC1 and DC2, is accompanied by three distinct waves of expression of transcription factors and signaling molecules that in concert are likely regulating the specification of the DC niche fate and initiation of DC niche function.

### DC Cell Fate Specification Requires Signals from Pre-existing Placodes

Our data suggest that the transition of fibroblasts to pre-DC for DC fate acquisition depends on the early wave of transcription factor and signaling molecule upregulation. However, it is unclear how the upregulation of these early DC genes is triggered, i.e. whether it is a pre-programmed cell-autonomous process or whether it relies on intercompartmental signals from epithelial placode progenitors. Understanding this developmental hierarchy essentially addresses the order of progenitor and niche fate specification. Previous studies have demonstrated that *Wnt/β-catenin* signaling in the epidermis is essential for placode formation and hair follicle morphogenesis including the formation of aggregated DCs (Huelsken et al., 2001; Zhang et al., 2008). In light of our identification of unclustered pre-DC cells, we asked whether an early DC fate is established in the absence of placode progenitors. To this end, we blocked placode formation by conditionally ablating *Wnt/β-catenin* signaling in the embryonic epidermis of *K14-Cre;β-catenin*^*flfl*^ mice. In E14.5 control skin we observed, as described above, multiple developmental stages of SOX2^+^ ITGA6^-^ DCs underneath EDAR^+^ placodes, including pre-DC (**Figures 6A** and **S6A**). Additional double immunofluorescence with FOXD1 confirmed pre-DC cell identity (**Figure S6B**). As predicted, in the *β-catenin*-ablated conditional knockout skin, EDAR^+^ placodes and established DC niches, aggregated SOX2^+^ ITGA6^-^ DC1 and DC2, were absent (**Figures 6B** and **S6A**). Intriguingly, we neither detected groups of unclustered SOX2^+^ ITGA6^-^ (**Figure 6B**) or SOX2^+^ FOXD1^+^ ITGA6^-^ (**Figure S6B**) pre-DC cells in the absence of placodes, indicating that DC cell fate acquisition was also blocked. These data demonstrate that DC cell fate acquisition from fibroblasts requires the presence of placode progenitor cells that likely provided essential placode-derived signals.

**Figure 6.**
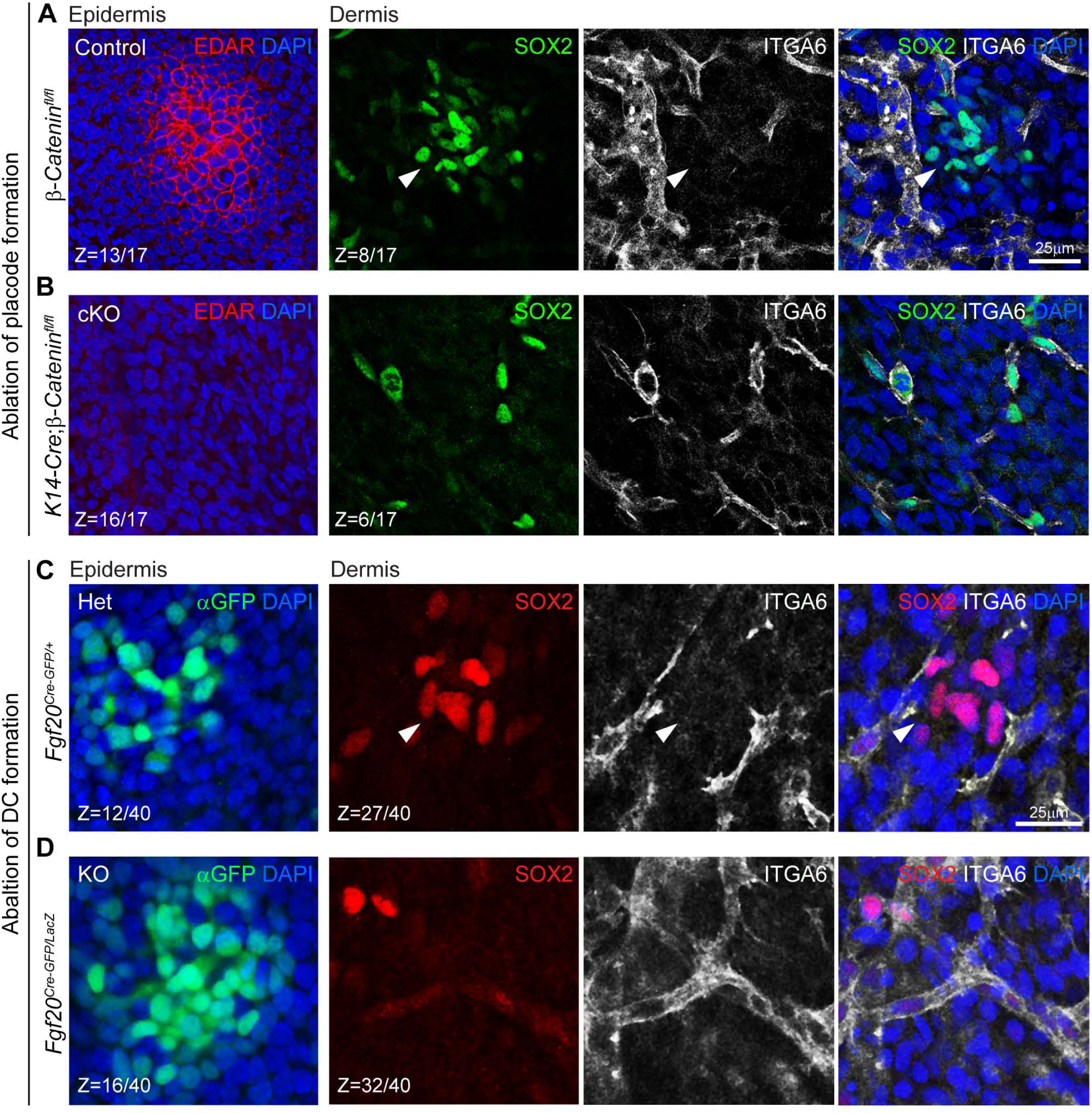
DC Cell Fate Specification Requires Pre-existing Placodes and Progenitor-Derived Fgf20. (A, B) Ablation of placode formation in *β-catenin* conditional knock out (cKO) epidermis. Immunofluorescence whole mount staining for EDAR, ITGA6 and SOX2. Z-sections from a 3D-confocal scan are shown at the epidermal and dermal level. Z values are section # of total optical sections in scan from epidermis into dermis. Section intervals = 1 μm. (A) Pre-DC (arrowhead) are specified underneath EDAR^+^ placodes in E14.5 *β-catenin*^*fl/fl*^ control embryos. (B) Pre-DC fail to become specified in *K14-Cre;β-catenin*^*fl/fl*^ cKO skins. (C, D) Ablation of DC fate specification in *Fgf20* knock out mice. Immunofluorescence whole mount staining for GFP, ITGA6 and SOX2. (C) Pre-DC (arrowhead) are specified underneath GFP^+^ placodes in E14.5 *Fgf20*^Cre-GFP/+^ control embryos. GFP is expressed in placode progenitors. (D) Pre-DC fail to become specified in *Fgf20*^Cre-GFP/LacZ^ KO knockout skins. Note Fgf20 promoter-driven GFP is still expressed in KO placodes.

### Placode Progenitor-Derived FGF20 Is Required For DC Cell Fate Specification

In searching for potential placode-derived signals essential for DC fate acquisition, we turned to *Fgf20*, a well-known downstream target of *Wnt/β-catenin* signaling that is required for DC formation (Huh et al., 2013). Recent studies have further demonstrated that FGF signaling is required for cell migration to form morphologically distinguishable DC clusters (Biggs et al., 2018; Glover et al., 2017). Having discovered pre-DC cells prior to DC formation, we asked whether *Fgf20* is required for DC niche fate specification before cell aggregation. To this end, we generated *Fgf20*-ablated embryos by crossing fertile and viable *Fgf20*^*Cre-GFP/Cre-GFP*^ knockout mice with *Fgf20*^*LacZ/+*^ mice; in both strains the first exon of the Fgf20 allele is replaced with a Cre-GFP fusion or LacZ gene (Huh et al., 2012, 2015). In E14.5 heterozygous *Fgf20*^*Cre-GFP/+*^ control skins, normal hair follicle development was observed with multiple DC stages underneath GFP^+^ placodes (Figures 6C and S6C). On the other hand, in Fgf20 knockout skins of *Fgf20*^*Cre-GFP/LacZ*^ embryos aggregated mature DCs were absent underneath GFP^+^ placodes, as described previously (Figures 6D and S6C) (Huh et al., 2013). Intriguingly, here like in the *Wnt/β-catenin* signaling ablation above, SOX2^+^/ITGA6^-^ or SOX2^+^/FOXD1^+^ pre-DC cells were also absent (**Figures 6C, S6C** and **S6D**), suggesting that FGF20 is a key placode-derived signal essential for DC fate specification, prior to cell aggregation.

## DISCUSSION

The timing of fibroblast-to-DC niche transformation has been traditionally described by the emergence of a characteristic tightly clustered DC beneath a thickened epidermal placode (Millar, 2002). It has been noted that at the earliest identified stage of *bona fide* HF morphogenesis, a distinct placode exists before the morphological DC cluster forms. Recently, two studies have demonstrated that the structured DC forms through directed migration of dermal fibroblasts (Biggs et al., 2018; Glover et al., 2017), but the precise timing and specific molecular drivers of DC fate-acquisition in relation to the physical aggregation process and to the timing of placode progenitor specification remained unexplored.

Here we demonstrate that *de novo* expression of the definitive (clustered) DC markers, *Sox2* (Clavel et al., 2012; Driskell et al., 2009; Sennett et al., 2015) and *Foxd1* (Sennett et al., 2015), first appears before any appreciable observed DC clustering. Taking advantage of *Sox2* and other DC molecular markers, both pre- and post-clustering, we were now able to define the molecular landscape of the earliest fate-acquired DC precursors and their most closely related fibroblasts, further refining our previous molecular characterization of the established DC niche (Sennett et al., 2015). Reinforcing our novel sorting strategy for population-based RNA sequencing analyses of distinct unclustered DC precursors and clustered DC stages, unbiased lineage prediction from single-cell transcriptomes not only reconstructed the DC differentiation path of the bulk approach but also provided a highly resolved account of the molecular transitional states and developmental trajectory from non-committed fibroblasts through DC2.

Complementing the model of DC niche formation through directed migration from *ex vivo* live imaging (Biggs et al., 2018; Glover et al., 2017), we found that while the fibroblasts surrounding the DC can have robust proliferative activity, the earliest fate-acquired, *Sox2*^+^ DC precursors have significantly diminished proliferation indicative that the mature DC is a product of continual fibroblast recruitment (**Figure 7A**). We interestingly observed an early steady decline in the expression of proliferation-related genes in fibroblasts most closely-related to pre-DC in pseudotime. This preemptive suppression of proliferation occurs prior to upregulation of early DC-fate genes and is particularly intriguing since the establishment of asymmetrically positioned DC are required in maintaining cell-fate asymmetry in the epithelium giving rise to both hair follicle stem cells and progenitors during early down growth of developing follicles (Cetera et al., 2018; Ouspenskaia et al., 2016; Xu et al., 2015). Therefore, by extension, it can be posited that tight regulation of DC size by proliferation suppression is vital to maintain its asymmetric positioning for proper morphogenesis and hair follicle stem cell establishment. The potential necessity of cell-cycle exit for acquisition of DC fate, the cell intrinsic and/or extrinsic mechanisms to achieve cell cycle arrest, and the precise molecular controls that prevent over-recruitment through migration remain to be studied in more detail.

**Figure 7.**
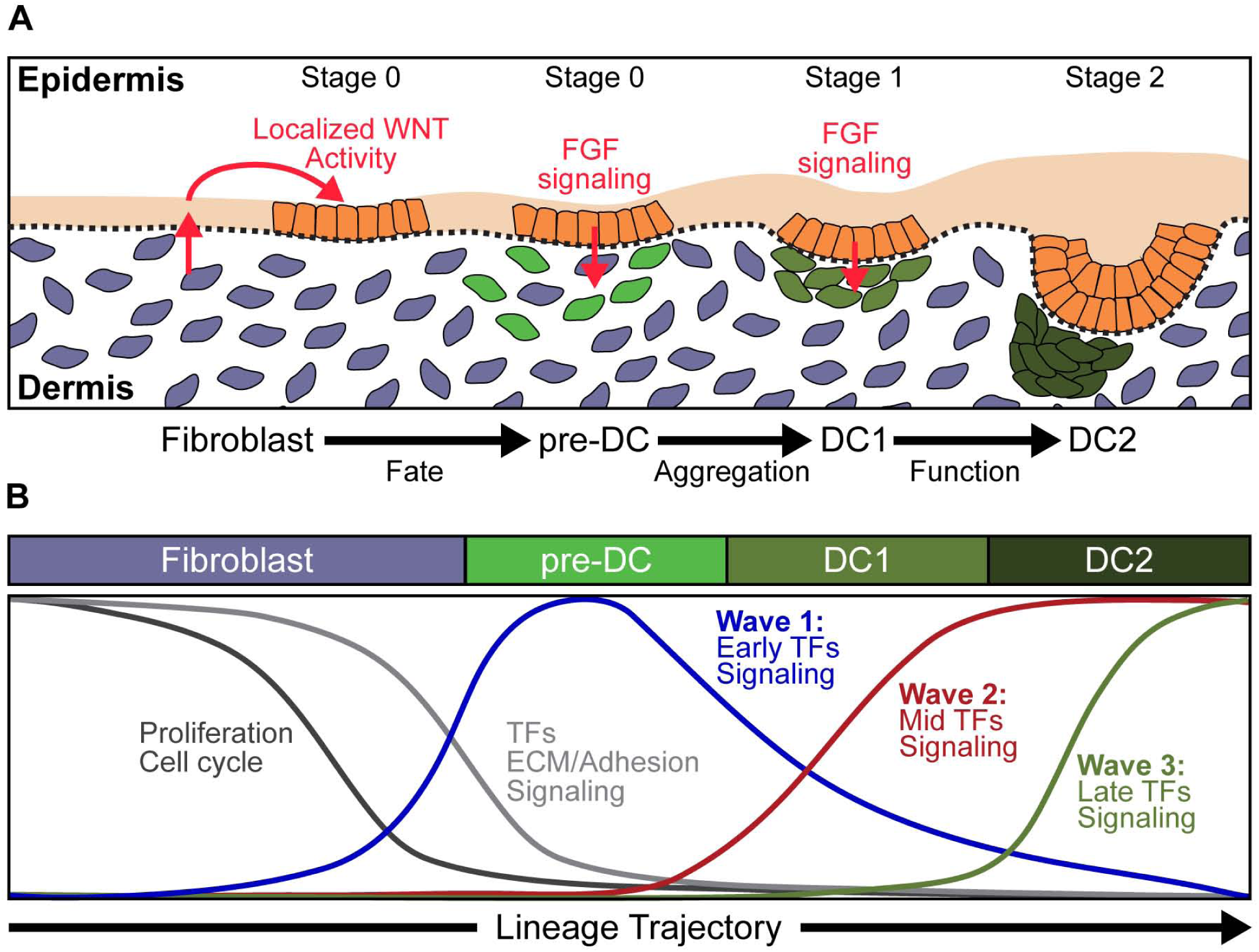
Model of DC Fate Specification and Maturation During Hair Follicle Formation. (A) Schematic depicting placode-derived FGF signaling-dependence of DC precursor fate specification before aggregation of the DC niche. (B) Key steps of the DC niche fate trajectory during the fibroblast-to-DC niche transition. Exit from the cell cycle and progressive loss of fibroblast fate coincide with progressive acquisition of the DC niche fate through three waves of DC transcription factors and signaling molecules expression.

From both the bulk and single cell transcriptome analyses, we identified transcription factors enriched in pre-DC including *Twist2, Lef1* and *Smad3* indicating an immediate shift in the gene regulatory network during early niche fate acquisition potentially acting as trailblazers for the downstream upregulation of mid- and late-expressed transcription factors and genes for DC formation and function such as those encoding cell-adhesion and signaling proteins (**Figure 7B**). The reconstructed lineage order in pseudotime also revealed a subset of fibroblasts closely related to pre-DC raising the possibility of a yet earlier precursor stage to the DC niche fate. While pinpointing their precise spatial location poses a considerable challenge because of the paucity of distinctly expressed genes to be used as molecular markers, clustering analyses place these cells broadly among upper dermal fibroblasts which harbor multipotent dermal fibroblast progenitors (Driskell et al., 2013).

Epithelial-mesenchymal interdependence during early HF morphogenesis has been shown by multiple compartment-specific disruptions affecting the development of its counterpart (Andl et al., 2002; Chen et al., 2012; Tsai et al., 2014; Zhang et al., 2009). By preventing placode formation through epidermal *β-catenin* ablation (Huelsken et al., 2001), we determined from the absence of pre-DC, early DC fate acquisition is not an autonomous pre-program of the upper dermis, but rather requires local directive signals from spatially patterned placodes. Genetic ablation of *Fgf20* has previously demonstrated its requirement for DC formation by hair follicle stage 1 through migration (Biggs et al., 2018; Huh et al., 2013), and furthermore here we show that FGF20 directly promotes the early DC niche fate. Similarly, recent identification of an Fgf20-expressing olfactory epithelial stem cell niche revealed a Wnt-regulated FGF20 requirement for formation of the underlying mesenchymal condensations that form nasal turbinates (Yang et al., 2018). Taken together these data indicate that the progenitors in nascent hair follicle pre-placodes signal to underlying fibroblasts at least in part through FGF20 to induce early DC fate prior to cluster-forming migration, and unequivocally demonstrate that progenitor fate precedes establishment of its supportive niche.

We now propose an update to the model of progenitor and niche specification during early hair follicle morphogenesis (**Figure 7A** and **7B**): Wnt-dependent placode progenitors signal to the underlying dermal fibroblasts via the FGF/FGFR signaling axis initiating a cascade of dynamic transcriptional waves that regulate DC niche fate specification. Through multifaceted signaling interactions between placode, pre-DC and unspecified interfollicular dermal fibroblasts, fate specified DC precursor cells centrally migrate to form the clustered DC (DC1). Continued pre-DC fate acquisition and migration incorporates additional cells to the DC cluster, while in the earliest established DC the new transcriptional regulatory network – set up by the first transcriptional wave – begins to upregulate expression of the molecular machinery crucial for signaling functions from the DC niche to the progenitors above.

## METHODS

### Mice

Specific strain and genotype details are provided in the Key Resources Table. Mice were housed in facilities operated by the Center for Comparative Medicine and Surgery (CCMS) at the Icahn School of Medicine at Mount Sinai (ISMMS) and all animal experiments were performed in accordance with the guidelines and approval of the Institutional Animal Care and Use Committee at ISMMS. Embryo ages were determined by designating the morning of the appearance of a vaginal plug as E0.5.

### EdU incorporation

For EdU-uptake proliferation assays, pregnant females mated with *Sox2*^*GFP*^ males were injected intraperitoneally with a single dose of EdU/PBS (500 μg/g body weight) 6, 12 or 24 hours prior to sacrifice. EdU detection was performed with the Click-It EdU AlexaFluor 647 Imaging Kit (Life Technologies) according to manufacturer’s instructions.

### Immunofluorescence staining

Tissue sections were embedded in OCT (Tissue Tek) and cut at a thickness of 10 μm with a cryostat (Leica). Whole-mounted embryo skins or sections from E15.0 embryos were fixed in 4% PFA and washed with PBS. Whole-mount samples were additionally permeabilized for 2 hr in 0.3% Triton X-100 in PBS. Tissues were blocked in PBS-Triton with BSA/NDS for 2 hour at room temperature. Primary antibody labelling against EDAR (goat, 1:100, R&D), SOX2 (Goat, 1:100, Stemgent), FOXD1 (Goat, 1:100, Santa Cruz), GFP (chicken, 1:800, Abcam), and CXCR4 (Rat, 1:100, BD Pharmagen) were carried out overnight at 4ºC. After PBS washes, samples were incubated with Rhodamine Red-X-, AlexaFluor 488-, AlexaFluor 555-, or AlexaFluor 647-conjugated donkey anti-goat, rabbit, chicken or rat secondary antibodies (Jackson Immunoresearch, Invitrogen). Samples were again PBS washed, counterstained with 4’6’-diamidino-2-phenilindole (DAPI) and mounted onto slides with antifade mounting medium.

### Image acquisition and processing

Whole mount immunofluorescence and stained tissue sections were imaged using Leica SP5 DMI confocal and Leica DM5500 widefield microscopes, respectively, both equipped with Leica LASAF software. For 3D whole mount imaging, z-stacks of up to 60 planes with 1 μm vertical intervals were acquired with Leica 40X, 63X or 100X oil-immersion lenses. 3D stack images from confocal microscopy were processed and analyzed with ImageJ/FIJI (NIH) or Imaris 3/4D image visualization and analysis software (Bitplane).

### RNA in situ hybridization

Skins were harvested from E15.0 embryos and were fixed in 4% PFA, embedded into OCT, and cut into 20μm-thick sections. Slides were dehydrated through a graded ethanol (EtOH) series (50%, 70% and 100% EtOH/H_2_O). Slides were stored at −20°C overnight in 100% EtOH. The following day, slides were air dried and fixed in 4% PFA for 15 minutes at room temperature. *In situ* hybridization for *Twist2* was performed with RNAscope ISH, according to the manufacturer’s instructions (RNAscope 2.5 HD Reagent Kit-RED, Advanced Cell Diagnostics). Following the RNAscope protocol, slides were processed for co-immunofluorescence and imaged as described above.

### E15.0 cell isolation and sorting

To generate single cell suspensions from E15.0 *Sox2*^*GFP*^ dorsal skins, tissues were collected and processed as previously described (Sennett et al., 2015). Briefly, back skins were harvested by microdissection and were pooled for FACS for bulk RNA-sequencing while skin from a single embryo was taken for single cell sorting prior to single cell RNA-sequencing. Skins were then digested in a dispase (Invitrogen)/ collagenase (0.03%, Worthington) solution with 20 U/µl of DNase (Roche) at 4º C overnight followed the next day by 30 minutes incubation at 37º C. Cells were then filtered through 40µm-pore nylon cell strainers, followed by centrifugation at 350xg for 10 minutes. Resuspended cell pellets were stained with primary antibodies against EpCAM (BrV605-conjugated rat Ab, 1:100, Biolegend), CXCR4 (PE-conjugated rat Ab, 1:100, Biolegend), ITAG6 (BrV421-conjugated rat Ab, 1:100, Biolegend), PDGFRA (biotinylated rat Ab, 1:50, Affymetrix). This was followed by secondary labeling with Streptavidin-APC (1:200, Biolegend). Dead cells were identified by DAPI uptake. Cells were sorted on a FACSAria II fluorescence cytometry sorter (BD Biosciences) for both bulk and single cell RNA-sequencing. For single cell RNA-sequencing, live fibroblasts, pre-DC, DC1, and DC2 were index sorted into 384-well hard shell plates (Biorad) pre-loaded with 5 μl of vapor-lock (QIAGEN) containing a 100-200 nl mix of RT primers, dNTPs, and synthetic mRNA spike-ins. After sorting, plates were immediately quick spun and frozen to ࢤ80°C.

### Real-Time qRT-PCR

Total RNA from the FACS-sorted cells was purified with the Absolutely RNA Nanoprep Kit (Agilent) and quantified with the NanoDrop spectrophotometer (Thermo Scientific). Reverse transcription was performed with the SuperScript III First-Strand Synthesis System using oligo(dT) primers (Invitrogen). Real-time qRT-PCR was performed with a LightCycler 480 (Roche) instrument using LightCycler DNA Master SYBR Green I reagents (Roche). Fold changes between samples were calculated based on the 2^-ΔΔCt^ method and normalized to *Gapdh*.

### cDNA library preparation for bulk RNA-sequencing

Total RNA from the FACS-sorted cells was purified with the Absolutely RNA Nanoprep Kit (Agilent). RNA concentration and quality were measured by the Agilent Bio-analyzer. Samples with RIN (RNA integrity number) scores of 9.6 or higher were further processed. 5 µl of RNA from each sample was used for reverse transcription, followed by amplification using the RNA Ovation RNA-Seq System V2 (NuGEN). Using the Ovation Ultralow DR Library System (NuGEN), cDNA libraries were generated from 100 ng amplified cDNA with a unique barcoded adaptor for each sample. Library concentration and quality were quantified by Qbit (Invitrogen) and the Agilent Bioanalyzer. Samples were then sequenced on the Illumina HiSeq 200 platform using a 50-nt single-read setting.

### cDNA library reparation for single cell RNA-sequencing

Single cell RNA-sequencing was done following the SORT-seq protocol (Muraro et al., 2016), which improves on the CEL-Seq2 protocol (Hashimshony et al., 2016) with automation. After sorting, cells were lysed at 65°C for 5 minutes. RT and second strand mixes were dispensed by the Nanodrop II liquid handling platform for cDNA library preparation (GC Biotech). For each 384 well plate, 384 primers (1 library of 384 cells) was used. Single cell double-stranded cDNAs were pooled and in-vitro transcribed for linear amplification. Illumina sequencing libraries were prepared using TruSeq small RNA primers (Illumina) and sequenced paired-end twice at a read length of 75 bp using the Illumina NextSeq.

### 4D time-lapse live imaging

*Tbx18*^*H2BGFP*^ E14.0 embryonic dorsal skin was harvested by microdissection in ice-cold PBS, sandwiched between a 8 µm Nuclepore filter (Whatman) and the bottom of a 35 mm Lumox culture dish (Greiner) such that the epidermal side of the skin was in contact with the Lumox membrane, i.e. the bottom of the dish. The skin-membrane sandwich was stabilized with Matrigel (Corning). Explants were cultured in DMEM (phenol red free, Invitrogen) containing 10% fetal bovine serum, 1% HEPES, and 1% penicillin-streptomycin. After overnight culturing, 3D Z-stack images of up to 20 planes with 2 μm vertical intervals were acquired at 10 or 25 minute intervals during a total of 16-20 hours using a Zeiss LSM880 equipped with a 20X Plan Apo 0.8 NA air objective or a Zeiss Axio Observer Z1 Yokogawa spinning disk equipped with a 20x Plan Apo 0.4 NA air objective. During imaging sessions, explants were maintained in a live-cell chamber at 37°C with 5% CO2 and humidity control.

### Quantification of DC clustering state

3D reconstructed images for nearest neighbor measurements of pre-DC, DC1, DC1 and interfollicular dermal fibroblasts were generated in Imaris. Automated segmentation of each cell was then performed by placing a point sphere at the geometric center of each nucleus from which each point’s discrete coordinates were used to quantify the absolute distances between nuclei. Multiple hypothesis testing of the data from the average distance between 5 nearest neighbors among cells from pre-DC, DC1, DC2 and interfollicular fibroblasts were performed using Anova: single factor.

### Quantification of proliferation by EdU-uptake

Percent EdU^+^ among pre-DC, DC1, DC2 and interfollicular dermal cells was manually counted from acquired 3D confocal scans in ImageJ/FIJI. *Sox2*GFP^+^/ITGA6^-^ cells were considered as DC cells whereas GFP^-^/ITGA6^-^ cells were counted as interfollicular dermal cells. For each injection time point skins from 2 embryos were analyzed, from which 3 DCs/regions of each morphological stage were quantified (i.e. 3 follicles x 4 stages per embryo per time point). Statistical analysis was performed using one-tailed student’s t-test.

### Quantification of proliferation by cell division during time-lapse live imaging

To measure proliferation with time-lapse live imaging, 4D images acquired from explants were processed and analyzed with Imaris. 15625um^2^ areas (350×350pixels) containing pre-DC or interfollicular dermis were obtained by cropping from larger 20X fields. Areas containing pre-DC were identified by locating DCs at the final time point and back tracing in time through the clustering process over a 6-hour period whereas the areas containing interfollicular dermis were defined as not containing any DC over the entire imaging session. Total cell numbers and mitotic events were quantified manually. Hypothesis testing was performed using student’s t-test.

### Bulk RNA-sequencing data analysiss

All raw RNA sequencing reads were mapped to the mouse genome (mm10) with TopHat v2.0.3 (Trapnell et al., 2009) coupled with the Bowtie2 (Langmead and Salzberg, 2012) and aligned with default parameters. Transcriptomes were assembled and fragments per kilobase per million reads (FPKM) for each gene were computed with Cufflinks v2.1.1 (Trapnell et al., 2010) with default parameters. Differentially expressed genes (DEGs) were identified using Cuffdiff (with default parameters except for the library normalization method was upper quartile normalization, where FPKMs were scaled via the ratio of the 75 quartile fragment counts to the average 75 quartile value across all libraries) and ANOVA with the Benjamini-Hochberg correction for multiple hypothesis testing with significance cut off FDR <0.05. The Fisher exact test was used for enrichment analysis with the same multiple hypotheses testing correction procedure and cut off. Principle component analyses (PCA) were performed for samples using BioJupies (Torre et al., 2018). Population signature genes were defined by DEGs with a FPKM≥1, and a fold enrichment≥2 compared to all other populations. Signature genes of multiple population overlaps featured in the Venn diagram analysis were defined similarly as individual-population signature genes by which each overlapping population must individually meet minimal expression level and fold-enrichment constraints.

Selection of genes used in developmental trajectory analyses were obtained by excluding genes with too low expression across all stages (FPKM_any population_ < 1) and genes with similar expression levels across each developmental stage (FPKM_max_ < 2*FPKM_min_). The remaining genes were log2 transformed and z-score standardized to obtain 0-mean and unit standard deviation. Gene clustering was performed with Morpheus (Broad Institute: https://software.braodinstitute.org/morpheus) using Spearman rank correlation with average linkage. Gene ontology and KEGG pathway enrichment analyses were carried out using Enrichr (Chen et al., 2013). Gene Set Enrichment Analysis (Subramanian et al., 2005) was performed using GenePattern (Reich et al., 2006).

### Single cell RNA-sequencing data analysis

Analysis of single-cell RNA-seq data was performed independently in an R environment (R 3.5.1, RStudio 1.1.453) and a Python environment (Python 3.6.5, Jupyter 1.0.0). Only samples with more than 6000 transcripts/cell were included in the analysis. Additionally, mitochondrial genes, ERCC spike-ins, and non-expressed genes were excluded from downstream analysis. The remaining 1383 cells and 18483 genes were analyzed independently using the RaceID3 package (R-based) (Grün et al., 2015, 2016; Herman et al., 2018) or the Scanpy package (Python-based) (Wolf et al., 2018).

For RaceID3 analyses, normalization was performed by simple rescaling, which is done by dividing the transcript counts per cell by the total number of transcripts, and multiplying this by the minimum total number of transcripts across cells. Principal components with overrepresentation of genes related to GO annotations “cell proliferation” and “cell cycle” were removed using the CCcorrect function. Evaluation of transcriptome similarities by 1-Pearson’s correlation coefficient followed by k-medoids clustering was used to cluster cells and t-distributed stochastic neighbor embedding (tSNE) (Van Der Maaten and Hinton, 2008) was used for visualization.

Using the Scanpy module, highly variable genes were extracted, data was log-normalized, and principal components were computed. Principal components showing significant overrepresentation of genes linked to the GO annotations “cell proliferation” and “cell cycle” in the top or bottom 1% quantile of loadings were removed in a fashion similar to the CCcorrect function in RaceID3. Cells were clustered using the Louvain algorithm (Blondel et al., 2008), and visualized using UMAP (McInnes and Healy, 2018).

Pseudotime analysis was performed on a subset of cells: clusters 4, 5, 6, 7, and 8 in RaceID3 analysis and clusters 1, 2, 3, and 5 in Scanpy analysis which comprise dermal condensate cells and fibroblast precursors of the latter. Cell order along pseudotime was inferred using StemID2 (Grün et al., 2016; Herman et al., 2018) or diffusion pseudotime (Haghverdi et al., 2016).

For easier reproducibility, the complete code is available online (see Key Resources Table and Data Resources).

Cumulative percent was calculated as the running sum of FPKM values divided by the total FPKM sum. Cumulative percent plots for early, mid, and late expressed genes were generated by calculating the mean cumulative percent values for each position (cell) in the SOM order.

### Software

RaceID3 and StemID2 were run on RStudio version 1.1.453. Scanpy was run in Python version 3.6.5 in a Jupyter Notebook version 1.0.0. The scripts run are available at https://github.com/rendllab and https://github.com/kasperlab.

### Data Resources

The accession number for the sequencing data reported in this paper is NCBI GEO:XXXX (bulk RNA-sequencing) and NCBI GEO:XXXX (single cell RNA-sequencing.)

## ACKNOWLEDGEMENTS

We thank Elena Ezhkova (ISMMS) for sharing mice and reagents. Many thanks to the personnel at the Flow Cytometry and Microscopy CoRE at ISMMS and to Venu Pothula for technical assistance during single cell sorting. Thanks also go to the personnel at the Genome Technology Center at NYU, and to Judith Vivié for expert assistance with single cell RNA sequencing at the Hubrecht Institute (Netherlands). Additionally, we thank Marja Mikkola (U. Helsinki) and Peggy Myung (Yale)) for helpful discussions and valuable comments. K.W.M was supported by The Science Appearance Career Development Award fellowship from the Dermatology Foundation. N.H. was supported by grants T32GM007280 from NIH/NIGMS and F30AR070639 from NIH/NIAMS. A.M. was supported by NIH grants U54-HL127624 (LINCS-DCIC) and U24-CA224260 (IDG-KMC). D.M.O. was supported by NIH grant HL111190 and R21DC017042. M.R. was supported by grants from NIH/NIAMS (R01AR071047; R01AR063151) and New York State Department of Health (NYSTEM-C029574), and a fellowship from the Irma T. Hirschl Trust.

## AUTHOR CONTRIBUTIONS

K.W.M., R.S., and M.R. conceived the overall project design. K.W.M., N.S., N.H., L.G., D.S., Y.S., and L.M.Y. performed experiments. K.W.M., N.S., N.H., and Z.W. analyzed bulk RNA-seq data. N.S., M.M., and T.J. performed bioinformatics analysis of single cell RNA-seq data. All authors discussed results and participated in the preparation and editing of the manuscript. K.W.M, N.S., N.H, and M.R. wrote the manuscript. M.R. supervised the study.

## Supplemental Information Inventory

## Supplemental Figures

- **Figure S1, Related to Figure 1** - Identification of Unclustered Dermal Condensate Precursors
- **Figure S2, Related to Figure 2** - Validation of Sorting Strategy and Population-Level Analysis of RNA-seq Data from Sorted DC Niche and Comparable Skin Populations
- **Figure S3, Related to Figure 3** - Resolving Developmental Trajectory with Louvain and RaceID3 Clustering
- **Figure S4, Related to Figure 4** - Fate-Acquired DC Precursors Demonstrate Rapid and Sustained Suppression of Proliferation
- **Figure S5, Related to Figure 5** - DC Fate Acquisition and Development is Marked by Three Discrete Waves of Transcription Factor and Signaling Molecule Upregulation
- **Figure S6, Related to Figure 6** - DC Cell Fate Specification Requires Pre-existing Placodes and Progenitor-Derived FGF20

## Supplemental Tables

- **Table S1, Related to Figure 2** – Gene Identities in Venn Diagram of Gene Expression Relationships Between Fb, pre-DC, DC1 and DC2
- **Table S2, Related to Figure 3** – Results of Hierarchical Clustering for Gene Trend Analysis
- **Table S3, Related to Figure 3** – Gene Identities in Self Organizing Map of Co-Regulated Gene Clusters Across Pseudotime
- **Table S4, Related to Figure 5** - Transcription Factors and Signaling Components Expressed During Early, Mid, and Late DC Gene Expression Waves

---

**Figure S1.**
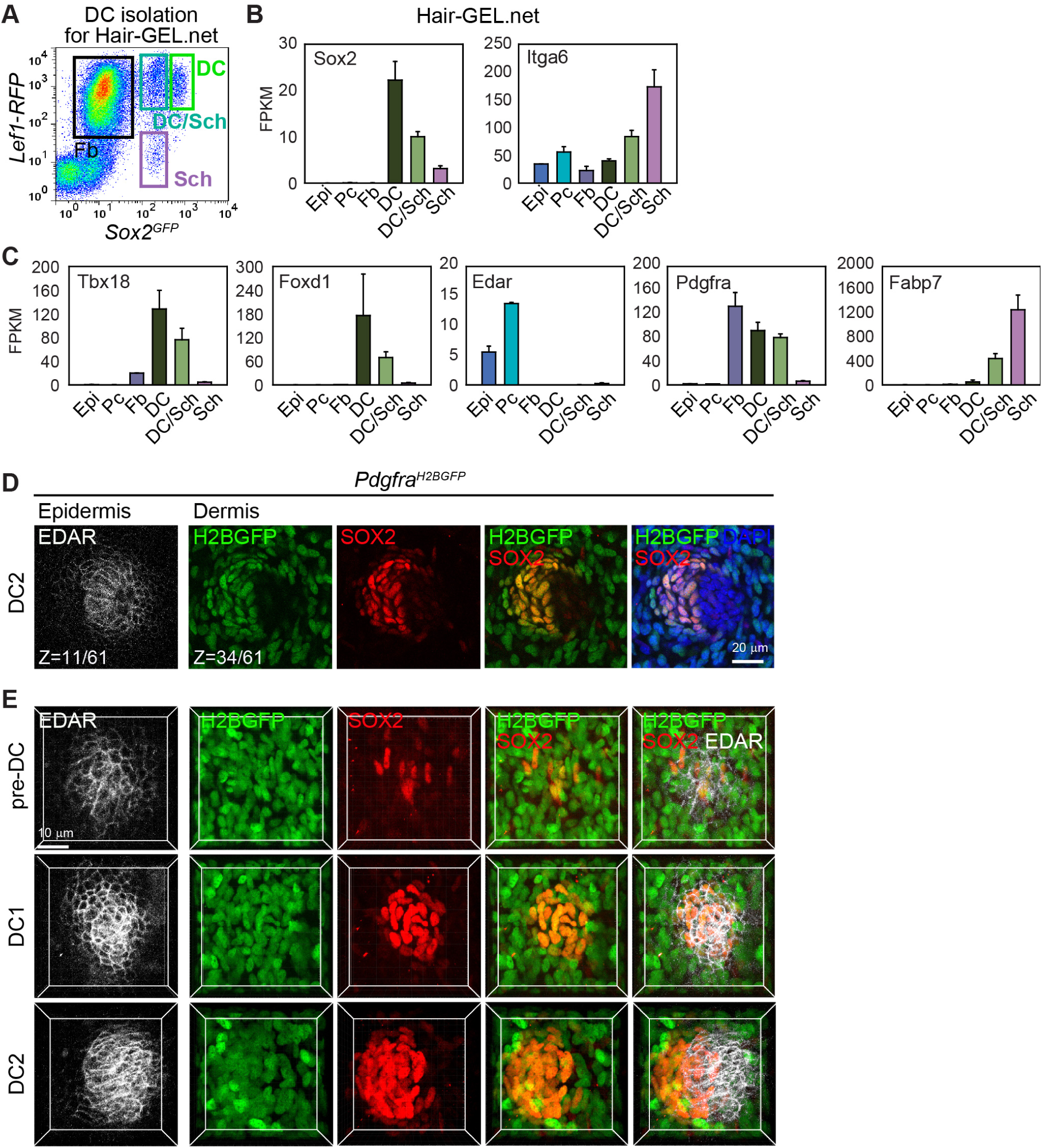
Related to Figure 1 – Identification of Unclustered Dermal Condensate Precursors. (A) FACS isolation of DC, Schwann cells (Sch), fibroblasts (Fb) and DC/Sch mix for RNA-seq from E14.5 *Sox2*^*GFP*^;*Lef1-RFP* back skin (Sennett et al., 2015). DC were sorted as GFP high, RFP^+^, Sch as GFP high, RFP ^*-*^, and DC/Sch mix as GFP low, RFP^+^. (B) Expression of *Sox2* (DC and Sch marker) and *Itga6* (Sch marker) in E14.5 sorted interfollicular epidermis (Epi), placode (Pc), Fb, DC, DC/Sch, and Sch. FPKM values are from Hair-GEL.net (Sennett et al., 2015). Data are mean ± SD. (C) Expression levels of additional known markers for DC (*Tbx18, Foxd1*), placode (*Edar*), Fb and DC (*Pdgfra*) and Sch (*Fabp7*). FPKM are from Hair-GEL.net (Sennett et al., 2015). Data are mean ± SD. (D) Immunofluorescence whole mount staining for EDAR and SOX2 on E15.0 *Pdgfra*^*H2BGFP*^ back skin. Nuclear H2BGFP marks mesenchymal cells. Z-sections from a 3D-confocal scan are shown at the epidermal and dermal level. Z values are section # of the total optical sections in a scan from epidermis to dermis. Section intervals = 1 μm. Note H2BGFP^+^/SOX2^+^ DC2 is asymmetrically positioned below the EDAR^+^ epidermal germ. (E) 3D rendered volumetric images from confocal scans of pre-DC, DC1 and DC2 stained with EDAR and SOX2 acquired from E15.0 *Pdgfra*^*H2BGFP*^ back skin were generated using Imaris image analysis software. Note H2BGFP^+^ SOX2^+^ nuclei of unclustered pre-DC cells lying directly underneath an early EDAR^+^ placode. Pre-DC and DC1 are the same as the Z-sections in Figure 1E.

**Figure S2.**
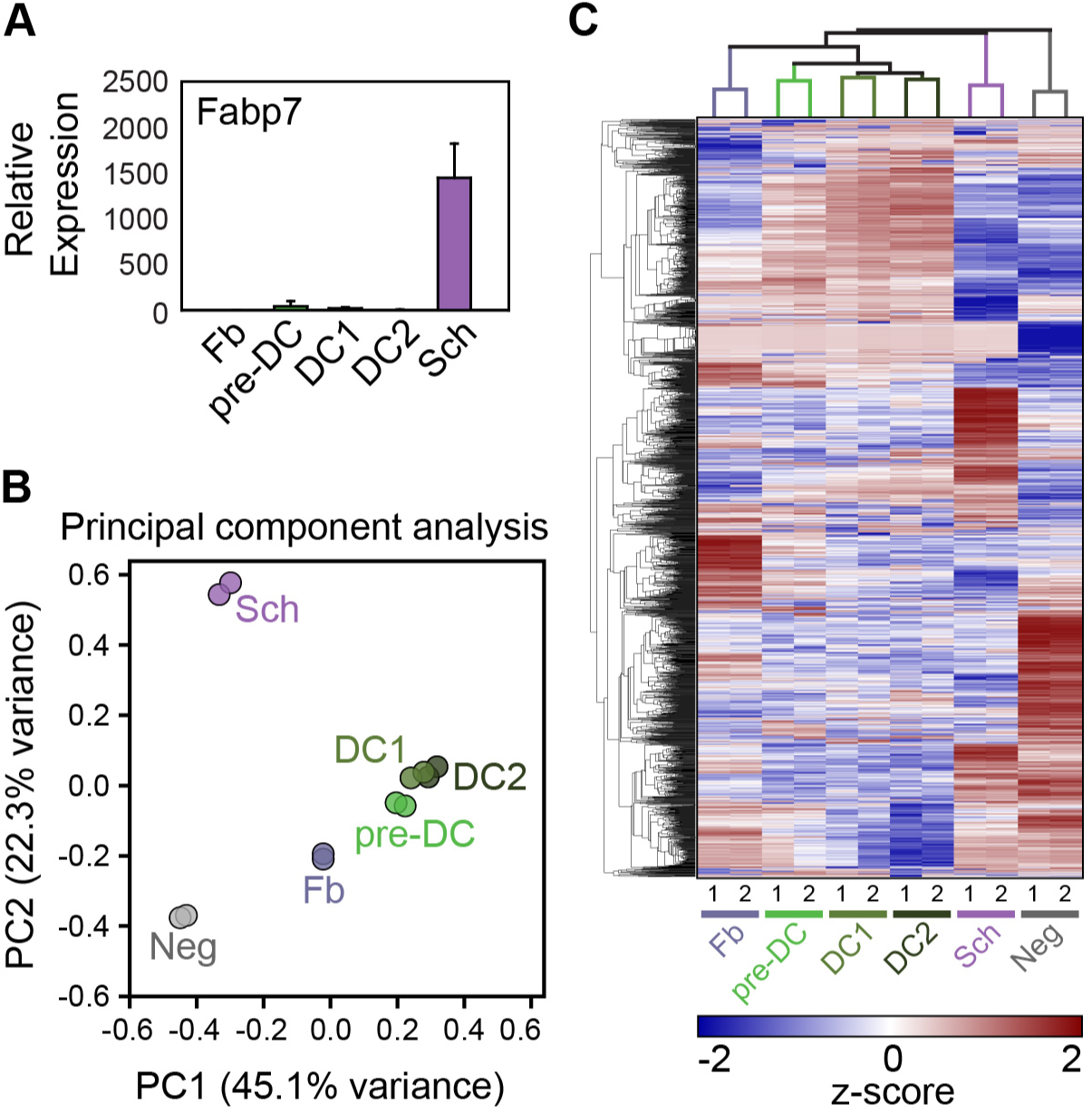
Related to Figure 2 – Validation of Sorting Strategy and Population-Level Analysis of RNA-seq Data from Sorted DC Niche and Comparable Skin Populations. (A) qRT-PCR of Sch marker *Fabp7* from FACS isolated Fb, pre-DC, DC1, DC2, and Sch. Data are mean ± SD. (B) Principal component analysis using the top 2,500 differentially expressed genes by ANOVA in Fb, pre-DC, DC1, DC2, Sch, and Neg cells. Note tight clustering of all DC populations and positioning of pre-DC between Fb and DC1/DC2. Each dot represents a single replicate. (C) Hierarchical clustering analysis using the top 2,500 differentially expressed genes by ANOVA. Euclidian distance clustering of populations (columns) and genes (rows). Z-scores were calculated from log_2_ transformed FPKMs and zero-centered.

**Figure S3.**
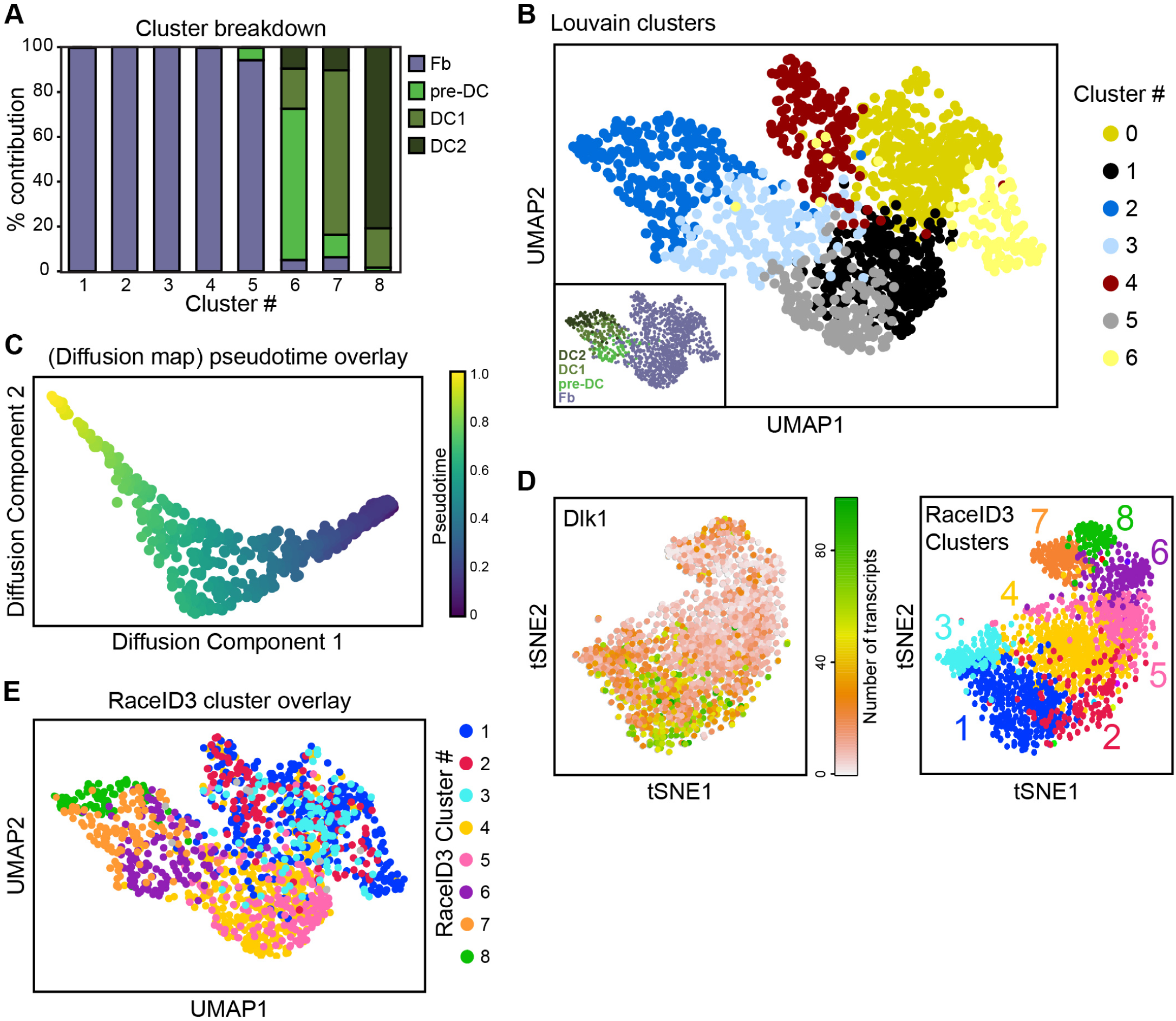
Related to Figure 3 – Resolving Developmental Trajectory with Louvain and RaceID3 Clustering. (A) Cell-type percent contribution breakdown of the 8 major RaceID3 clusters. (B) UMAP plot of Louvain clustering (Scanpy) of Fb, pre-DC, DC1, and DC2. Inset: UMAP plot of Louvain clustering with sorted cell identities highlighted from Figure 3C. (C) Diffusion map with pseudotime overlay for Louvain clusters 1, 2, 3, and 5 from (B). (D) Left: tSNE plot of *Dlk1* in RaceID3 clustering of Fb, pre-DC, DC1, and DC2. Right plot: RaceID3 clusters for reference from Figure 3A. Note highest *Dlk1*-expressing cells in RaceID3 output Fb clusters 1, 2 and 3. (E) UMAP plot from Louvain clustering colored to identify RaceID3 clusters.

**Figure S4.**
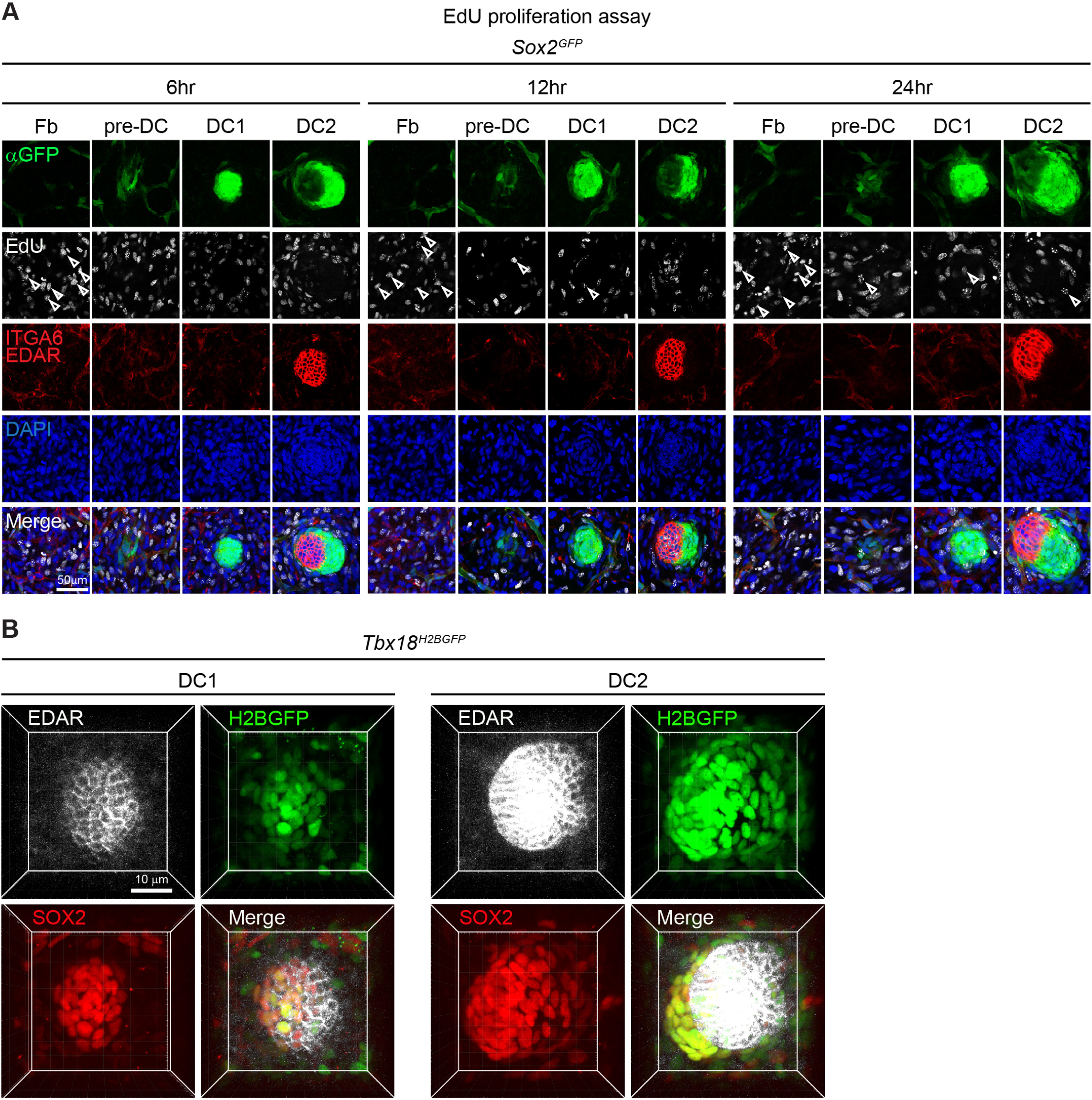
Related to Figure 4 – Fate-Acquired DC Precursors Demonstrate Rapid and Sustained Suppression of Proliferation. (A) Individual channels of GFP, EdU, ITGA6/EDAR, and DAPI from merged images Figure 4E. (B) 3D volume rendering from confocal scans of DC1 and DC2 stained for EDAR and SOX2 from E15.0 *Tbx18*^*H2BGFP*^ back skin were generated using Imaris. Note H2BGFP^+^ SOX2^+^ nuclei of DC1 lying directly underneath an early EDAR^+^ placode whereas those of DC2 lie in an anterior-posterior polarized position relative to the placode.

**Figure S5.**
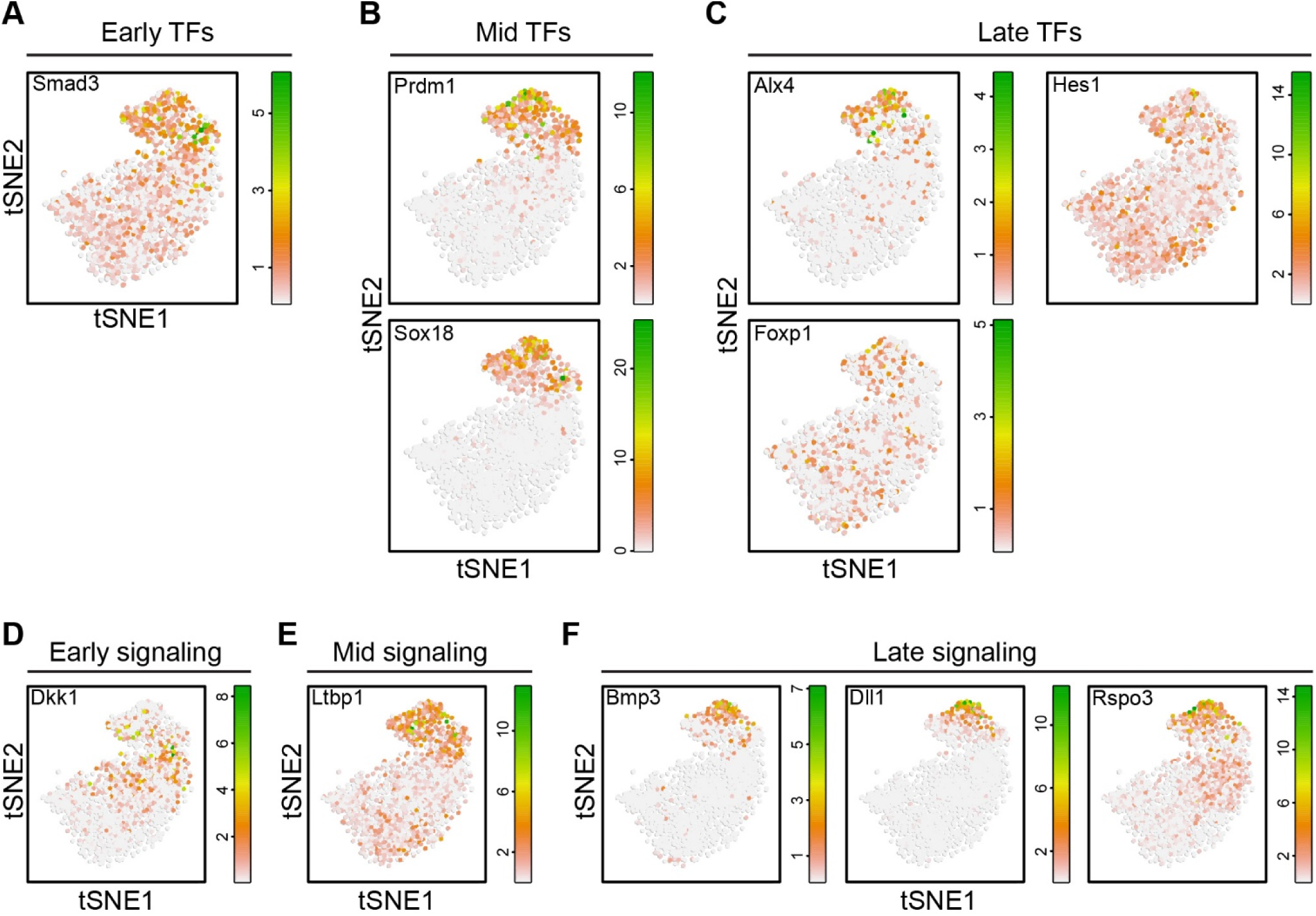
Related to Figure 5 – DC Fate Acquisition and Development is Marked by Three Discrete Waves of Transcription Factor and Signaling Molecule Upregulation. (A) tSNE plot of early-expressed transcription factor *Smad3*. (B) tSNE plots of mid-expressed transcription factors *Prdm1* and *Sox18*. (C) tSNE plots of late-expressed transcription factors *Alx4, Foxp1* and *Hes1*. (D) tSNE plot of early-expressed signaling molecule *Dkk1*. (E) tSNE plot of mid-expressed signaling molecule *Ltbp1*. (F) tSNE plots of late-expressed signaling molecules *Bmp3, Dll1,* and *Rspo3*.

**Figure S6.**
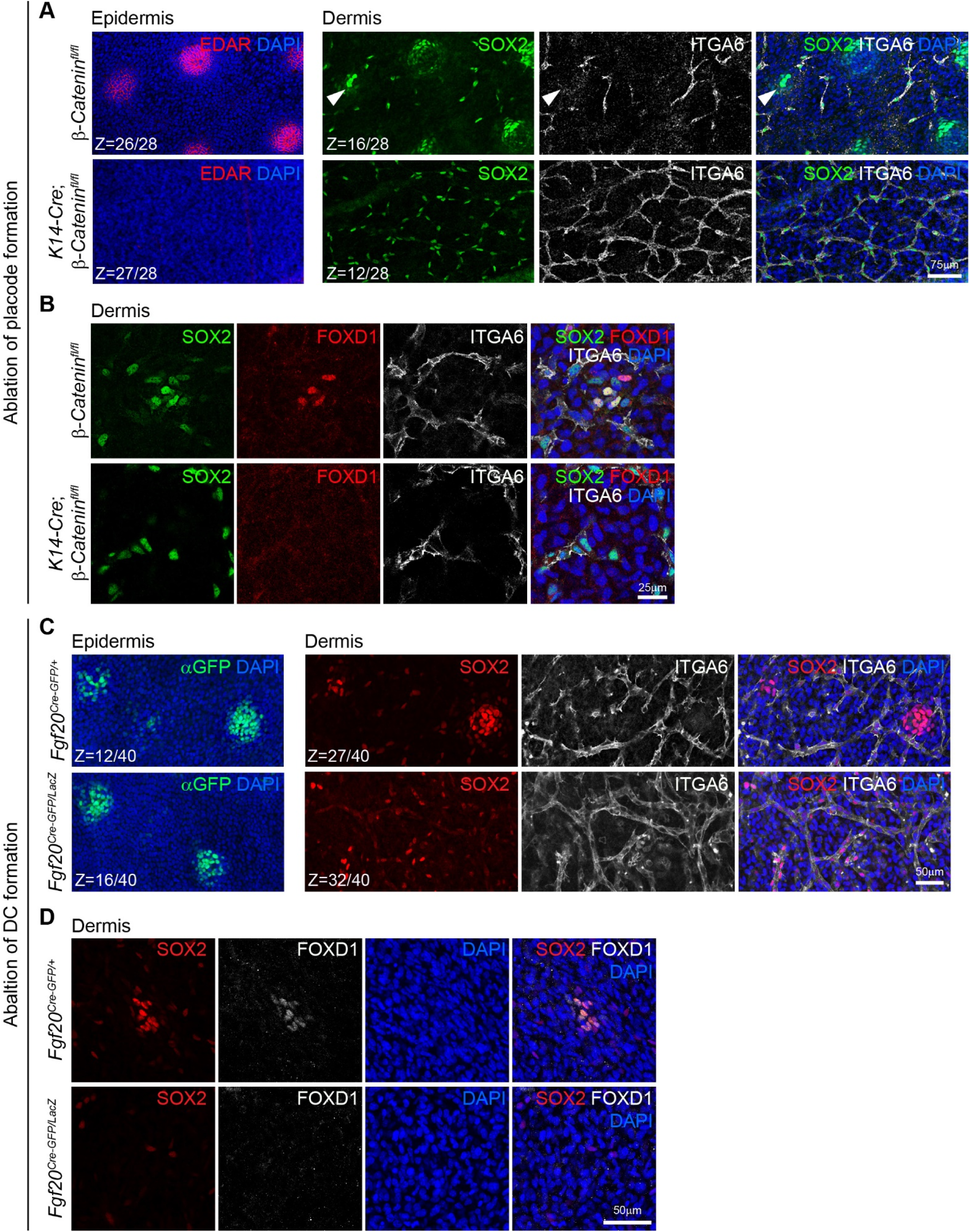
Related to Figure 6 – DC Cell Fate Specification Requires Pre-existing Placodes and Progenitor-Derived FGF20. (A) Low magnification confocal optical sections of whole mount immunofluorescence for EDAR, SOX2, and ITGA6 of E14.5 control (upper panels, β-*catenin*^*fl/fl*^) and conditional epidermal knockout (lower panels, *K14-Cre;*β-*catenin*^*fl/fl*^) skin. Left panels: optical sections at level of epidermis show EDAR^+^ placodes in control but no EDAR-expressing cells in mutant epidermis. Right panels: sections at dermal levels show presence of SOX2^+^ ITGA6^-^ pre-DC (arrowhead), clustered DC1 and DC2 below hair placodes in control dermis, but no SOX2^+^ ITGA6^-^ cells are observed in mutant dermis. (B) Optical section of E14.5 control and *K14-Cre;*β-*catenin*^*fl/fl*^ whole mount stained back skins for SOX2, FOXD1, and ITGA6. Note FOXD1^+^ pre-DC in control skin and absence of any FOXD1^+^ dermal cells in the knockout. (C) Low magnification confocal optical sections of whole mount immunofluorescence for GFP, SOX2, and ITGA6 of E15.0 control (upper panels, *Fgf20*^*Cre-GFP/+*^) and knockout (lower panels, *Fgf20*^*Cre-GFP/LacZ*^) skin. Left panels: optical sections at level of epidermis show GFP^+^ placodes in both control and mutant epidermis. Right panels: sections at dermal levels show SOX2^+^ ITGA6^-^ pre-DC (arrowhead), clustered DC1 and DC2 below hair placodes in control dermis, but no SOX2^+^ ITGA6^-^ pre-DC, DC or DC2 are observed in mutant dermis. (D) Optical section of E15.0 control and *Fgf20*^*Cre-GFP/LacZ*^ whole mount stained back skins for SOX2 and FOXD1. Note FOXD1^+^ pre-DC in control skin and absence of any FOXD1^+^ dermal cells in the knockout.

**Table S4.**
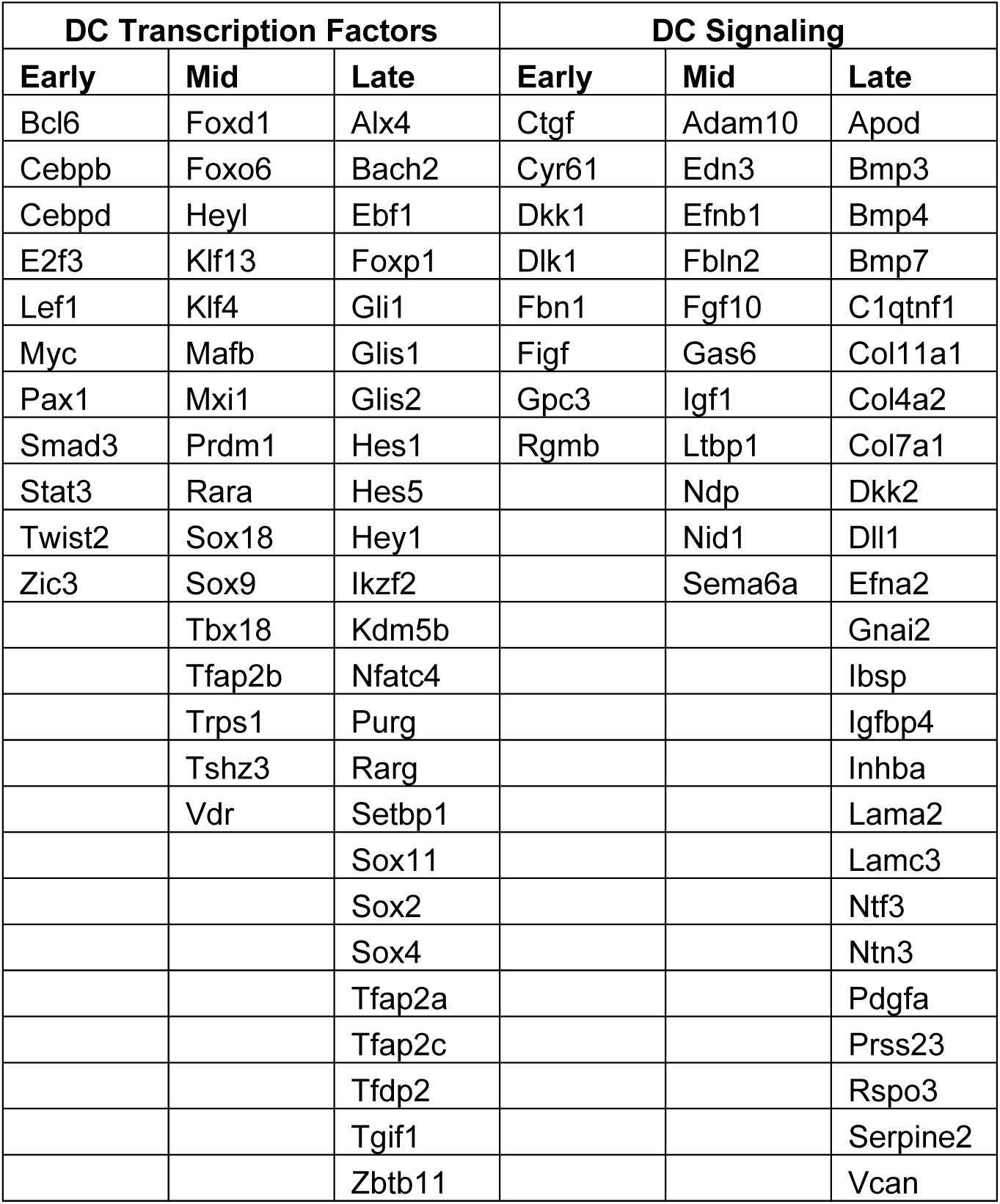
Related to Figure 5 – Transcription Factors and Signaling Components Expressed During Early, Mid, and Late Gene Expression Waves

